# Inducible Hepatocyte Aryl Hydrocarbon Receptor Ablation Prevented TCDD-induced Steatosis and Fibrosis but Enhanced Hyperplasia and Hypertrophy

**DOI:** 10.64898/2026.11.26.769307

**Authors:** Karina Orlowska, Rance Nault, Giovan N Cholico, Zachary Smith, Warren Sink, Anna Wojtowicz, Alex Vien, Fiona Utley, David Vinnikov, Candice Robillard, Nathan Shami, Jennifer Robinson, Jordan Stubleski, Rachel H. Wilson, Christopher A. Bradfield, Jack Harkema, Tim Zacharewski

## Abstract

The aryl hydrocarbon receptor (AHR) mediates 2,3,7,8-tetrachlorodibenzo-*p*-dioxin (TCDD)-induced hepatotoxicity, including steatosis, inflammation, and fibrosis. To investigate the role of hepatocyte AHR we used an inducible hepatocyte-specific AHR knockout model (Ahr^dV375Afl/fl^;Alb-CreERT2). Male control and knockout mice received 0-30 µg/kg TCDD every four days for 28 days. Hepatocyte-specific AHR ablation was confirmed by loss of TCDD-induced CYP1A1 in hepatocytes with evidence of preserved induction in non-parenchymal cells. Hepatocyte AHR ablation prevented TCDD-induced steatosis and attenuated lipid peroxidation, bile acid accumulation, and fibrosis. Although immune cell infiltration increased in both genotypes, induction of macrophage-associated genes was lower in knockout mice, whereas inflammatory cytokines and chemokines remained elevated. Unexpectedly, TCDD treated knockout mice exhibited marked hepatocellular hypertrophy, and hyperplasia. These findings demonstrate that AHR expression in hepatocytes is required for steatosis, macrophage remodeling, and fibrotic progression with roles in regulating hepatocyte proliferation.

## Introduction

The aryl hydrocarbon receptor (AHR) is a ligand-activated transcription factor that plays a central role in mediating cellular responses to a wide range of environmental, dietary, and endogenous chemicals, yet no universally accepted cognate ligand has been identified (*1*). In its inactive state, AHR resides in the cytosol as part of a complex with other chaperone proteins. Upon ligand binding, the complex translocates into the nucleus, where the AHR dissociates from chaperone proteins and heterodimerizes with the AHR nuclear translocator (ARNT). The AHR-ARNT complex then binds to dioxin response elements (DREs) to regulate gene expression involved in xenobiotic metabolism, cell proliferation, inflammation, and metabolic homeostasis. AHR-mediated gene expression can also occur through DRE-independent mechanisms following interaction with other transcription factors (*2*). Although the canonical AHR signaling pathway is best known for driving the induction of phase I and II metabolizing enzymes, accumulating evidence indicates that the AHR also serves broader physiological functions, including the modulation of liver development, regeneration, and energy balance (*3*).

2,3,7,8-Tetrachlorodibenzo-*p*-dioxin (TCDD) is the prototypical high-affinity AHR ligand that elicits species-, sex-, age-, tissue-, and cell-specific changes in gene expression (*1*). In the liver, TCDD elicits characteristic, AHR-dependent pathologies that mirror the hepatic features of metabolic dysfunction-associated steatotic liver disease (MASLD), namely the progression of steatosis to steatohepatitis with fibrosis. Within hours of TCDD treatment, AHR target genes are dose dependently induced including cytochrome P450s and reactive oxygen species (ROS) defenses with evidence of lipid accumulation that resolves after the cessation of exposure due to the short half-life of TCDD in mice (8-12 days) compared to humans (1-11 years) (*4–6*). However, following repeated oral gavage, the severity of the hepatic phenotype intensifies with more pronounced steatosis accompanied by inflammation, hepatocyte hypertrophy, biliary epithelial proliferation, and fibrosis (*7*).

The AHR exhibits ubiquitous cell and tissue distribution with the highest expression in hepatocytes based on normalized transcripts per million according to the Human Protein Atlas (proteinatlas.org) (*8*). Several mouse models have been developed to investigate the physiological and toxicological significance of the AHR. Independent knockout studies have demonstrated the importance of AHR in normal liver development and in mediating the toxic effects of environmental contaminants (*9*) (*10*) (*11*). Although *Ahr*^−/−^ mice were viable, they exhibited developmental liver abnormalities, including delayed growth, reduced liver weight, transient microvesicular steatosis, prolonged extramedullary hematopoiesis, and portal hypercellularity accompanied by fibrosis (*9*). These models also established that AHR is essential for the toxic responses to TCDD with *Ahr*^−/−^ mice being largely resistant to up to 2,000 µg/kg TCDD, a dose ∼10-fold higher than that causing severe toxicity in mice expressing functional AHR (*10*). Moreover, TCDD-induced effects in the liver, thymus, and other organs were largely absent in AHR-deficient mice. Subsequent studies further demonstrated that AHR deficiency prevented other TCDD-induced changes in organ weights, hepatic gene expression, and bile acid metabolism (*12*). Similar findings were reported for another AHR ligand, benzo[a]pyrene (BaP), as AHR deficiency prevented BaP-induced *Cyp1a1* expression and skin tumor formation (*13*).

Despite these advances, early AHR knockout models had limitations. First, constitutive AHR deletion during development resulted in hepatic abnormalities that could influence subsequent responses, making it difficult to distinguish the direct effects of AHR loss in adult tissues from developmental consequences. Second, differences in AHR ligand-binding affinity among mouse strains substantially affect their responsiveness to TCDD. The low-affinity *Ahrᵈ* allele, found in DBA/2 mice and incorporated into some conditional knockout models, exhibits ∼10-fold lower binding affinity for TCDD compared to the high-affinity *Ahrᵇ* allele expressed in C57BL/6 mice (*14*). Consequently, mice expressing the *Ahr*^d^ allele require significantly higher doses for many AHR ligands, which increases the possibility of off-target effects. More specifically, the ED_50_ for TCDD-induced hepatic *Cyp1a1* mRNA induction was 0.08 µg/kg in C57BL/6J mice compared with 1.5 µg/kg in DBA/2J mice (*15*). Similarly, LD_50_ values of 159 and 3,351 µg/kg reported in congenic C57BL/6J mice expressing the *Ahrᵇ* and *Ahrᵈ* alleles, respectively, corresponding to a ∼21-fold difference in susceptibility to TCDD-induced lethality (*16*). Mice with the *Ahrᵈ* allele also required an 8 to 24-fold higher dose of TCDD to induce hepatocellular changes, increase liver weight, and alter serum biomarker levels (*16*). To address these issues, a conditional floxed AHR mouse model (*Ahr^dfl/fl^*) was generated, enabling cell-specific deletion of AHR when crossed with mice expressing Cre recombinase under tissue-specific promoters (*17*). However, this model was derived from 129-SvJ embryonic stem cells that expressed the low-affinity *Ahrᵈ* allele, with lower binding to TCDD and other ligands. Consequently, the ligand-binding domain in the *Ahr^dfl/fl^*model was modified using CRISPR by substituting the valine at position 375 with alanine, generating a conditional AHR model that harbors high-affinity allele, *Ahr^d^*^V375Afl/fl^ (*18*). This modification enabled tissue-specific investigation of AHR function using TCDD doses more consistent with those used in C57BL/6 mouse studies. Furthermore, no studies have examined the dose dependent hepatic effects of TCDD in an *in vivo* model where the AHR has been deleted in hepatocytes following normal liver development.

In the present study, *Ahr^dV375Afl/fl^* females were crossed with male *Alb-CreERT2* mice to obtain a tamoxifen (TAM)-inducible, hepatocyte-specific AHR knockout mouse model (*Ahr^dV375Afl/fl^*;Alb*-CreERT2* hereafter referred to as AHR^HEP^) that offers temporal control over *Ahr^dV375A^*deletion. Following normal liver development, AHR^HEP^ mice were treated with TAM to generate AHR knockout mice to investigate the role of AHR expression in hepatocytes in mediating the dose-dependent effects of TCDD while preserving AHR signaling in non-parenchymal liver cells. Following oral gavage of TCDD every 4 days for 28 days, control (AHR^HEP^-CTRL) and knockout (AHR^HEP^-TAM) mice were examined for effects on liver weight, serum biomarkers, histopathology, lipid accumulation, inflammation, fibrosis, hepatocyte hypertrophy and hyperplasia and differential gene expression. The results revealed distinct TCDD induced AHR-dependent and -independent responses and provided further evidence of AHR regulating hepatocyte hypertrophy and proliferation.

## RESULTS

### Confirmation of hepatocyte-specific AHR ablation

In this study, a TAM-inducible hepatocyte-specific *Ahr* knockout mouse model (AHR^HEP^-TAM) was used that expresses a modified *Ahr^d^* allele with high binding affinity for TCDD (*18*). This model provided temporal control and targeting of AHR^dV375Afl/fl^ deletion in hepatocytes. Untreated AHR^HEP^ (*Ahr^dV375Afl/fl^*;Alb*-CreERT2*) mice exhibited no overt morphological or behavioral abnormalities. Developmental milestones, food and water intake, reproductive performance and sex ratio in untreated AHR^HEP^ mice were comparable to wild type C57BL/6Ncrl mice.

At approximately post-natal day (PND) 32, male and female AHR^HEP^ mice were intraperitoneally injected with either corn oil (AHR^HEP^*-*CTRL) or 75 mg/kg TAM (AHR^HEP^*-*TAM) for five consecutive days followed by a five day clearance period (*19*). Body weight gain was lower in TAM treated AHR^HEP^ mice. TAM is reported to induce physiological stress in mice that can reduce body weight gain (*20–22*). To confirm *Ahr^dV375A^* deletion, mice received a single oral gavage of 30 µg/kg TCDD for 24 hrs (**Figure 1A**). PCR analysis showed that TAM produced the expected shorter transcript, whereas AHR^HEP^-CTRL mice retained exon 2 with negligible excision of *Ahr^dV375A^* likely due to CreERT2 “leakiness” in the absence of TAM (**Figure 1B**). To confirm hepatocyte loss of AHR^dV375A^ protein, primary hepatocytes were isolated from both AHR^HEP^*-*CTRL and AHR^HEP^*-*TAM mice and treated with TCDD for 24 hours. TCDD dose-dependently induced *Cyp1a1* in primary hepatocytes isolated from AHR^HEP^*-*CTRL mice, whereas no induction was detected in AHR^HEP^*-*TAM hepatocytes, consistent with the AHR-mediated induction of CYP1A1 following binding to dioxin response elements (**Figure 1C**). Furthermore, immunohistochemistry detected strong TCDD-induced CYP1A1 expression in AHR^HEP^*-*CTRL liver sections (**Figure 1D-2**) with negligible hepatocyte staining in AHR^HEP^*-*TAM liver sections (**Figure 1D-3**). However, CYP1A1 staining was still evident in non-parenchymal cells (**Figure 1D-3 inserts**) in AHR^HEP^*-*TAM liver sections. Together, these results confirm that TAM treatment caused robust and selective ablation of AHR in hepatocytes but not in non-parenchymal cells.

**Figure 1.**
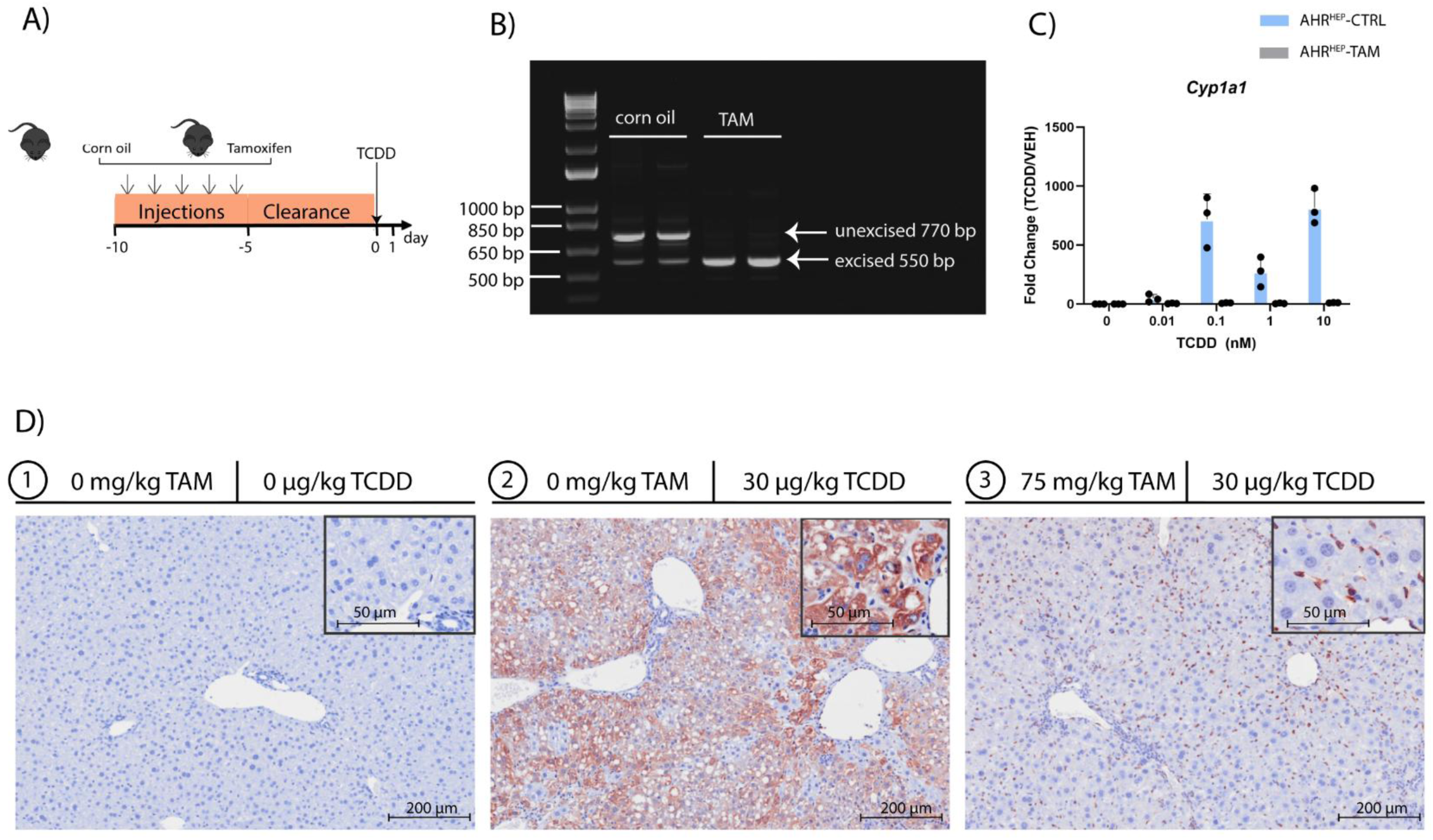
Tamoxifen (TAM)-induced hepatocyte-specific AHR ablation in male AHR^HEP^ mice. **A)** Experimental design. *Ahr* ablation was achieved in hepatocytes following intraperitoneal injection of *Ahr^V375A-fl/fl^*;*Alb-CreERT2* (AHR^HEP^) mice with 75 mg/kg tamoxifen for five consecutive days. TAM was then allowed to clear for 5 days prior to oral gavage with 30 µg/kg TCDD. **B)** Polymerase chain reaction (PCR) of *Ahr* shows exon 2 excision in AHR^HEP^-TAM mouse livers with negligible excision in AHR^HEP^-CTRL samples. **C)** Quantitative PCR (qPCR) of *Cyp1a1* in primary hepatocytes isolated from AHR^HEP^-CTRL and AHR^HEP^-TAM mice. Hepatocytes were treated with DMSO (vehicle) or 0.001 – 10 TCDD nM for 24 hours (n=3). **D)** Immunohistochemistry of CYP1A1 revealed cell specific staining in AHR^HEP^-CTRL and AHR^HEP^-TAM liver sections collected 24 hours after a single oral gavage with 30 µg/kg TCDD. The top right inserts at higher magnifications show widespread CYP1A1 staining in AHR^HEP^-CTRL hepatocytes as well as other cell types. In contrast, AHR^HEP^-TAM sections show CYP1A1 staining in non-parenchymal cells with negligible staining in hepatocytes.

**Figure 2.**
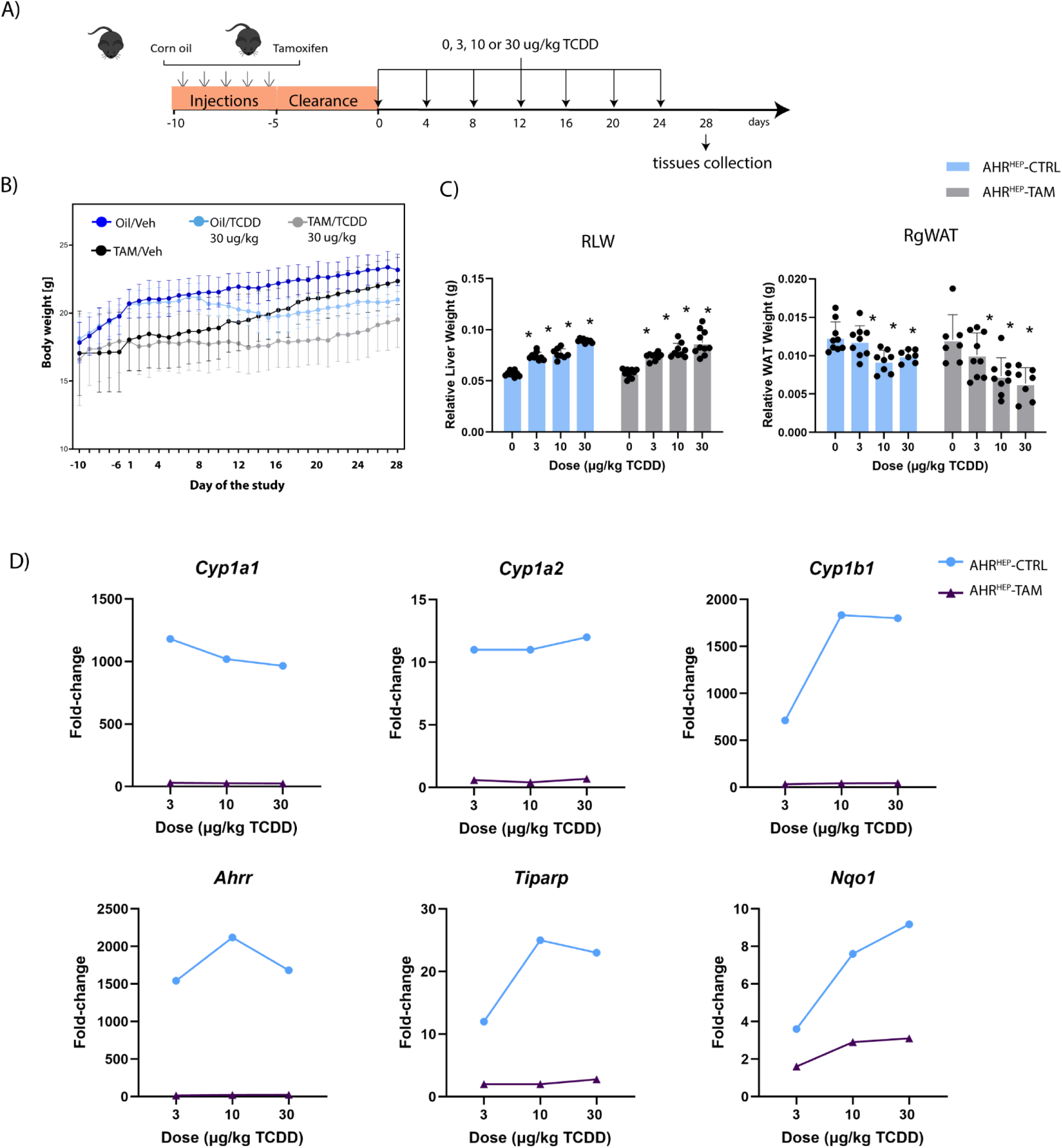
Experimental design and responses to TCDD in male AHR^HEP^-CTRL and AHR^HEP^-TAM mice. **A)** Schematic of experimental design. Hepatocyte-specific AHR ablation was achieved by intraperitoneal injection of *Ahr^V375A-fl/fl^*;*Alb-CreERT2* (AHR^HEP^) mice with 75 mg/kg TAM for five consecutive days, followed by 5 recovery days to allow TAM clearance. Mice were then orally gavaged with sesame oil vehicle or TCDD every 4 day for 28 days. **B)** Body weight assessment. **C)** Relative liver weight (RLW) and relative gonadal white adipose tissue (RgWAT) weight at day 28, 4 days after the last gavage. Bars represent mean ± SD. Asterisk (*) indicates significant difference (p < 0.05) determined by two-way ANOVA and Tukey’s post-hoc test (n=6). **D)** Differential expression of AHR battery genes in AHR^HEP^-CTRL and AHR^HEP^-TAM mice following oral gavage with TCDD every 4 days for 28 days. Gene expression was assessed by RNA-Seq (n=5).

**Figure 3.**
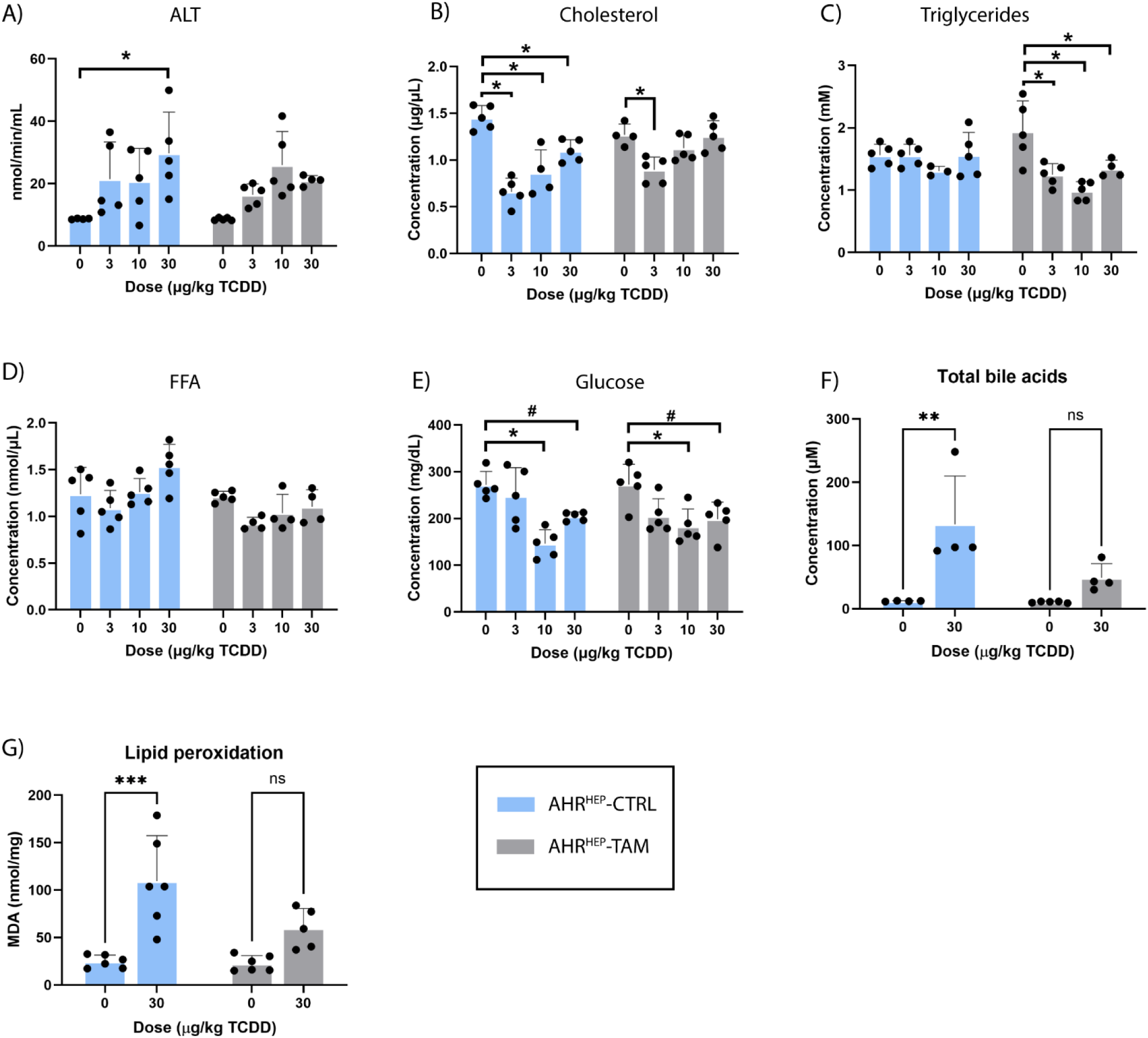
Effect of TCDD on serum clinical chemistry, serum bile acids level and hepatic lipid peroxidation in male AHR^HEP^-CTRL and AHR^HEP^-TAM mice. A) Serum alanine aminotransferase (ALT), B) total cholesterol, C) triglycerides, D) free fatty acids (FFA), and E) glucose in AHR^HEP^*-*CTRL and AHR^HEP^*-*TAM male mice orally gavaged every 4 days for 28 days with vehicle (sesame oil), 3, 10, or 30 µg/kg TCDD. F) Serum total bile acid levels and G) hepatic malondialdehyde (MDA) levels in AHR^HEP^*-*CTRL and AHR^HEP^*-*TAM male mice orally gavaged every 4 days for 28 days with vehicle (sesame oil) or 30 µg/kg TCDD. Bars represent mean ± SD. Asterisk (*p ≤ 0.05, **p < 0.01, ***p ≤ 0.001, ****p ≤ 0.0001) and pound sign (#) (p < 0.1) indicates significant difference determined by two-way ANOVA and Tukey’s post-hoc test (n=5).

### Physiological responses to TCDD

At PND 42–44, male and female AHR^HEP^*-*CTRL and AHR^HEP^*-*TAM mice were orally gavaged with vehicle (sesame oil), 3, 10 or 30 µg/kg TCDD every 4 days for 28 days to assess the effects of hepatocyte-specific AHR deletion (**Figure 2A, Supp. Fig. S1A**). AHR^HEP^*-*TAM male mice experienced less body weight gain compared to AHR^HEP^*-*CTRL mice with some recovery by day 28 (**Figure 2B**). TCDD treated mice experienced reduced body weight gain in both models that did not fully recovery by day 28 but was more severe in AHR^HEP^*-*TAM mice. Overall, AHR knockout in hepatocytes did not appreciably affect the repression of body weight gain by TCDD. There were negligible TAM- or TCDD-related effects on body weight gain in female mice (**Supp. Fig. S1B**) and no apparent difference in chow consumption (**Supp. Fig. S1C**). TCDD also dose-dependently increased relative liver weight (RLW) in both male and female models that was accompanied by a decrease in a relative gonadal white adipose tissue (RgWAT) weight (**Figure 2C**, **Supp. Fig. S1D-E**).

We then examined the expression of known hepatic AHR target genes following oral gavage with TCDD every 4 days for 28 days. As expected, TCDD induced expression of AHR-battery genes was dramatically greater in AHR^HEP^-CTRL mice but not completely abolished in AHR^HEP^-TAM mice (**Figure 2D**). For example, *Cyp1a1*, *Ahrr* and *Tiparp* were induced ∼1000-, ∼2000- and ∼24-fold in AHR^HEP^-CTRL mice but only ∼25-, 24- and ∼2-fold in AHR^HEP^-TAM mice, respectively. This is consistent with the preserved expression of AHR^dV375A^ in non-parenchymal cells in AHR^HEP^-TAM mice (**Figure 1D**) (*23, 24*). Furthermore, the TCDD induced levels of *Cyp1a1*, *Cyp1a2*, *Cyp1b1*, *Ahrr*, *Tipart* and *Nqo1* were comparable between AHR^HEP^-CTRL and wild type C57BL/6Ncrl mice (GSE203302) despite some evidence of leaky CreERT2 activity **(Figure 1B)**.

The dose dependent effects of TCDD on serum biomarkers were also examined (**Figure 3**). ALT levels (**Figure 3A**) were modestly increased while total cholesterol levels (**Figure 3B**) decreased in AHR^HEP^-CTRL mice. Triglycerides were reduced in AHR^HEP^-TAM mice by TCDD (**Figure 3C**) whereas free fatty acids were unaffected in both models (**Figure 3D**). Glucose levels were comparably reduced by TCDD in AHR^HEP^-CTRL and AHR^HEP^-TAM mice consistent with the induction of *Hk1*, *2* and *3* in both models with no AHR genomic binding 2 hrs after oral gavage with TCDD in wild type C57BL/6Ncrl mice (**Figure 3E, Supp. Fig. S3**). Collectively, these results suggest that increases in ALT and decreases in total cholesterol and serum glucose levels occurred independent of AHR expression in hepatocytes.

TCDD increased bile acid levels in AHR^HEP^-CTRL mice, as previously reported in wild-type C57BL/6Ncrl mice but did not significantly (p =0.48) induce bile acid levels in AHR^HEP^-TAM mice suggesting the effects of TCDD on bile acid levels were dependent on AHR expression in hepatocytes (**Figure 3F**) (*25*). Furthermore, TCDD induced malondialdehyde (MDA) levels in AHR^HEP^-CTRL mice whereas the increase in AHR^HEP^-TAM mice was non-significant (p > 0.05) (**Figure 3G**). This was consistent with the AHR-mediated induction of ROS producers (*Cyp1a1*, *Cyp1a2*, *Por*, *Xdh*) and ROS amplifiers (*Pdk3*, *Pdk4*) as well as the induction of *Ucp2*, an adaptive response to excess ROS, all of which exhibited greater induction in AHR^HEP^-CTRL mice (**Supp. Fig. S2**). Accordingly, several indicators of cellular stress or injury were more highly induced in AHR^HEP^-CTRL mice compared to AHR^HEP^-TAM mice such as *Cacng1*, *Cyfip2*, *Ecel1*, *Insl6*, *Pkp1*, *Spire2*, *Trpv4*, and *Krt25*. ROS defense responses were also more highly induced in AHR^HEP^-CTRL mice compared to AHR^HEP^-TAM mice including *Nfe2l2*, *Nqo1*, *Akr1b7*, *Aldh3a1*, *Aldh3b1*, *Slc7a11*, *Gsta1*, *Gpx2*, *Ugt1a7c*, *Ugt1a8* and *Ugt1a9* (**Supp. Fig. S2**). Many of these genes are regulated by AHR and/or NFE2L2 (*26–28*). Like the effects on bile acid, TCDD induced lipid peroxidation appears to be dependent on AHR expression in hepatocytes.

### TCDD Tissue Levels

TCDD levels exhibited dose dependent increases in the liver, gWAT and serum. However, the levels differed significantly within and between AHR^HEP^-CTRL and AHR^HEP^-TAM tissues (**Figure 4, Supp.Table S1**). The TCDD levels in male AHR^HEP^-CTRL liver samples were comparable to hepatic levels reported in similarly treated wild type female C57BL/6Ncrl hepatic samples (*29*). TCDD is sequestered in the liver following hepatic fat accumulation and the AHR-mediated induction of CYP1A2 **(Figure 2B)** (*30, 31*). Consequently, hepatic TCDD levels were significantly greater in AHR^HEP^-CTRL liver samples compared to AHR^HEP^-TAM mice. In contrast, serum and gWAT levels were significantly greater in AHR^HEP^-TAM mice suggesting that TCDD that was normally sequestered in the liver was redistributed to other tissues due to the lack of hepatocyte CYP1A2 induction and hepatic fat accumulation. For example, TCDD levels were ∼8-fold greater in AHR^HEP^-CTRL liver samples while serum and gWAT levels were both ∼4-fold greater in AHR^HEP^-TAM compared to AHR^HEP^-CTRL samples after oral gavage with 30 µg/kg TCDD every 4 days for 28 days. This also suggests extrahepatic tissues would likely experience greater TCDD exposure. The effects of AHR^dV375A^ loss on the elimination of TCDD via urine and feces were not examined.

**Figure 4.**
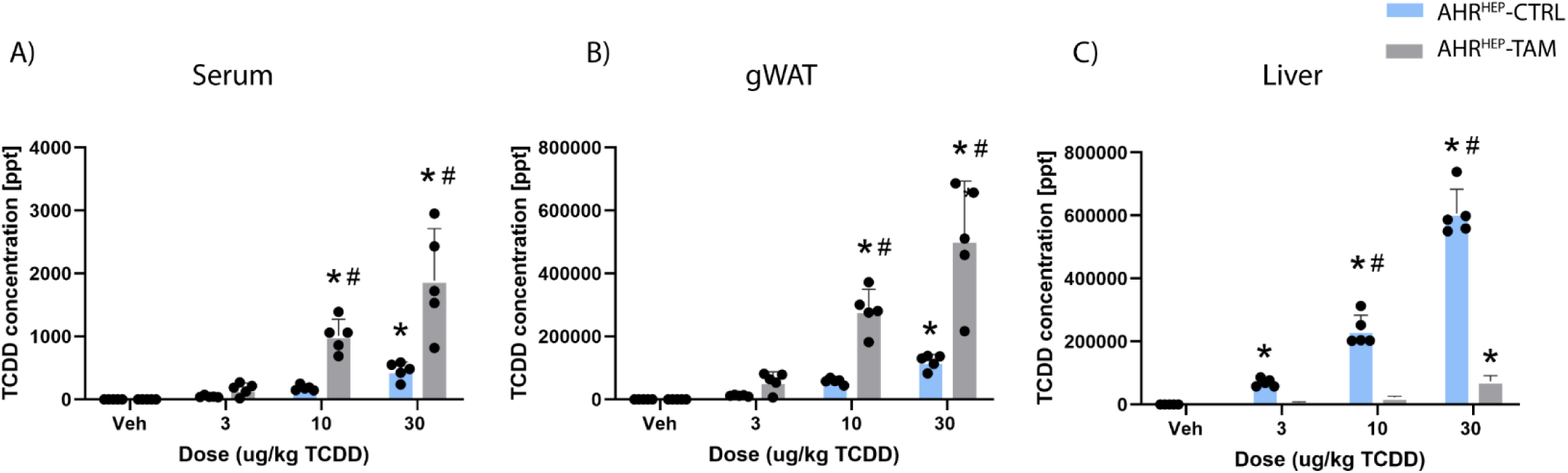
Dose-dependent accumulation of TCDD in serum, gonadal white adipose tissue (gWAT), and liver of AHR^HEP^-CTRL and AHR^HEP^-TAM male mice. TCDD concentrations in A) serum, B) gWAT, and C) liver of control (corn oil, CTRL) or tamoxifen (TAM)-treated AHR^HEP^ male mice orally gavaged every 4 days for 28 days with vehicle (sesame oil), 3, 10, or 30 µg/kg TCDD. TCDD concentrations are expressed in parts per trillion (ppt). Bars represent mean ± SD (n = 5 per group). Asterisk (*) indicates difference (p < 0.05) between TCDD and sesame oil vehicle within the same genotype and pound sign (#) indicates significance difference (p < 0.05) between AHR^HEP^-CTRL and AHR^HEP^-TAM for the same TCDD dose determined by two-way ANOVA followed by Tukey’s post-hoc test.

### Gross Morphology & Histopathology

TCDD decreased body weight gain without affecting food intake, as previously reported (**Supp. Fig. S1B, C**). However, TAM and TCDD co-treatment reduced body weight gain more than TCDD alone. **Table 1** summarizes the H&E assessment of liver sections following oral gavage with TCDD every 4 days for 28 days. H&E staining of control (vehicle) AHR^HEP^*-*CTRL and wild type C57BL/6Ncrl liver sections were comparable with no remarkable histological differences despite negligible evidence of CreERT2 leakiness in the absence of TAM. TCDD dose dependently induced the progression of steatosis to steatohepatitis with fibrosis in AHR^HEP^*-*CTRL, similar to reports in wild type C57BL/6Ncrl mice (*25, 32*). There was also evidence of mild to moderate apoptosis/necrosis and biliary hyperplasia as well as hepatocellular hypertrophy and hyperplasia in AHR^HEP^*-*CTRL, as previously reported in wild type C57BL/6Ncrl mice (*25, 32*). Inflammation apoptosis/necrosis, vacuolization and fibrosis were more severe in AHR^HEP^*-*CTRL mice. In contrast, AHR^HEP^*-*TAM mice had substantially greater hepatocellular hyperplasia and hypertrophy with evidence of hepatocytes in various stages of mitosis (**Table 1**, **Figure 6C**). Although there was reduced CYP450 induction (**Figure 2D**), lack of hepatic fat accumulation based on Oil Red O (**Figure 5B**), and a non-significant increase in lipid peroxidation (**Figure 3G**) with negligible induction of cellular stress or injury, AHR^HEP^*-*TAM mice exhibited mild to moderate hepatic inflammation (**Table 1**).

**Figure 5.**
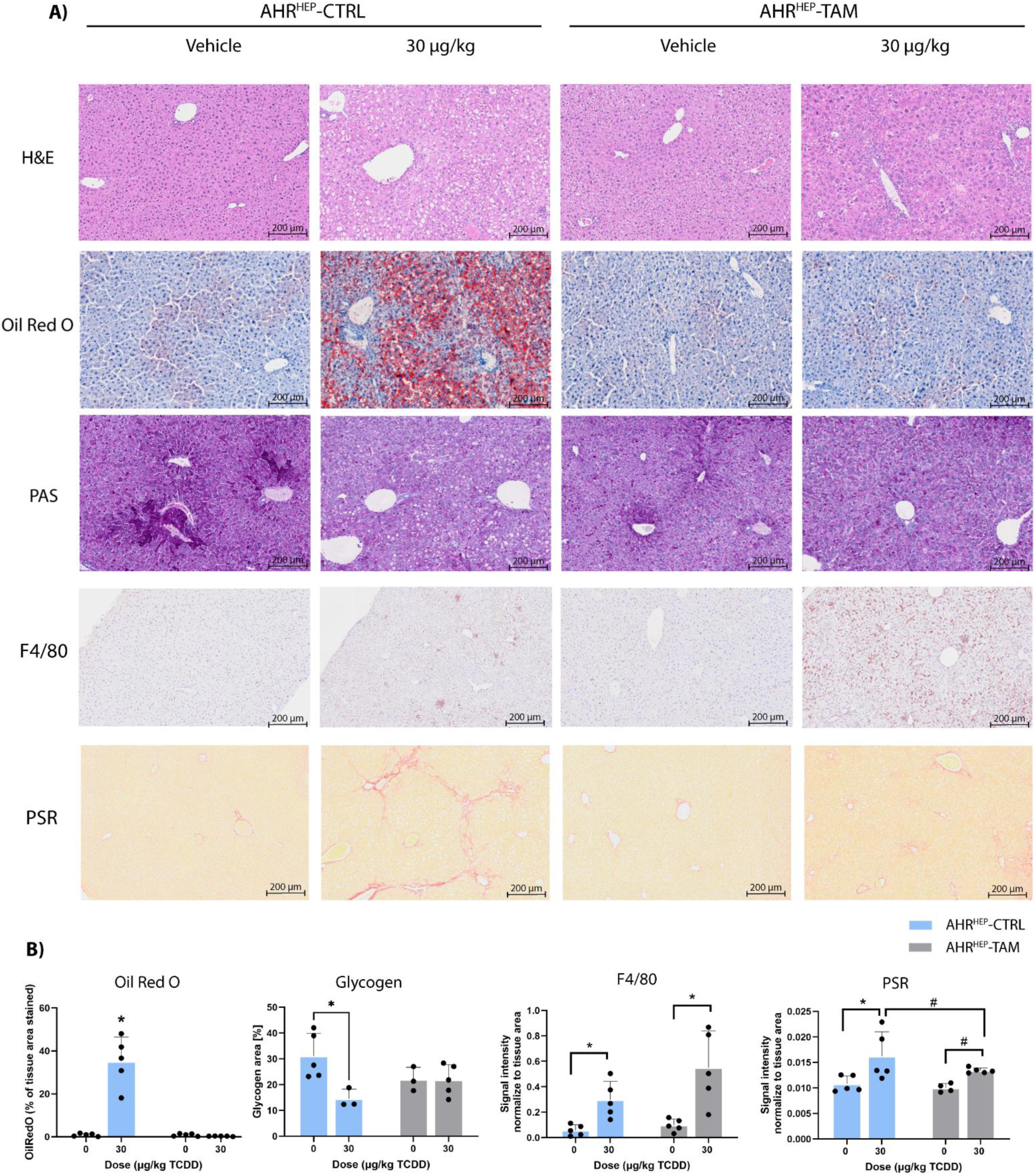
Representative micrographs of TCDD elicited hepatic effects. AHR^HEP^*-*CTRL and AHR^HEP^*-*TAM gavaged with vehicle (sesame oil) or 30 μg/kg TCDD every 4 days for 28 days**. A)** representative hematoxylin and eosin (H&E), Oil Red O (ORO), Periodic acid–Schiff (PAS), F4/80 and Picro-Sirius Red (PSR) stained liver sections. Scale bar represents 200 μm. **B)** ORO, PAS, F4/80 and PSR staining were quantified using QuPath software. Data is presented as mean ± SD (n=5). Asterisk (*) (p < 0.05) and pound sign (#) (p < 0.1) indicate significance difference as determined by two-way ANOVA and Tukey’s post-hoc test.

**Figure 6.**
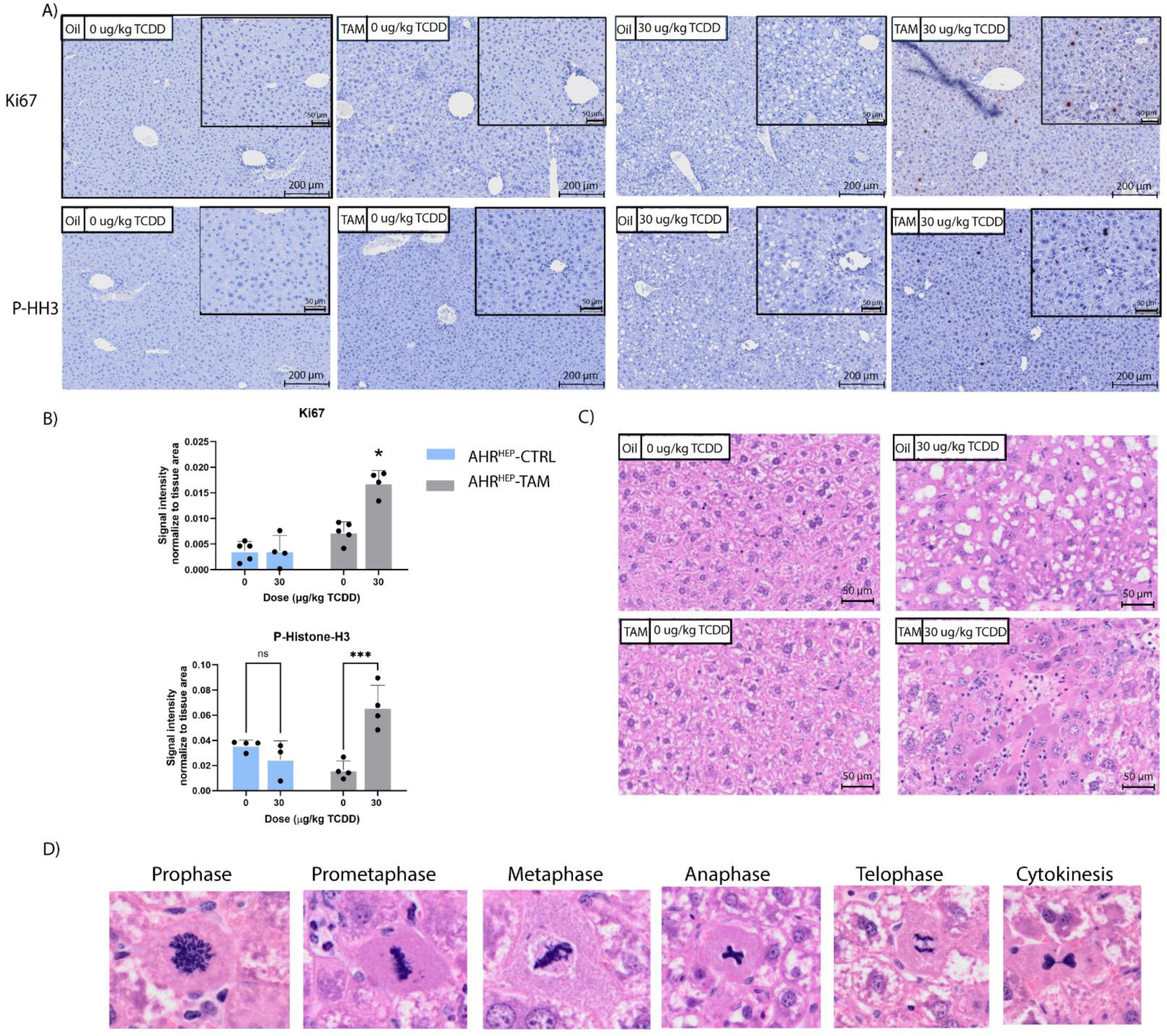
Hyperplasia and hypertrophy of hepatocytes in TCDD treated AHR^HEP^*-*TAM liver. AHR^HEP^*-*CTRL and AHR^HEP^*-*TAM mice were orally gavaged with vehicle (sesame oil) or 30 μg/kg TCDD every 4 days for 28 days**. A)** Ki67 and phospho-histone-H3 (P-HH3) staining in liver sections from AHR^HEP^*-*CTRL and AHR^HEP^*-*TAM mice treated with TCDD. Higher magnification is presented in top right insert. **B)** Quantitation of Ki67 and P-HH3 stained cells using the QuPath software. Data is presented as mean ± SD (n=4 or 5). Asterisk indicates significance (*p ≤ 0.05, ***p ≤ 0.001) determined by two-way ANOVA and Tukey’s post-hoc test. **C)** Representative H&E-stained sections from AHR^HEP^*-*CTRL and AHR^HEP^*-*TAM following oral gavage with 30 μg/kg TCDD every 4 days for 28 days. **D)** Representative mitotic figure images from AHR^HEP^*-*TAM H&E-stained liver sections following oral gavage with 30 μg/kg TCDD every 4 days for 28 days.

**Table 1.**
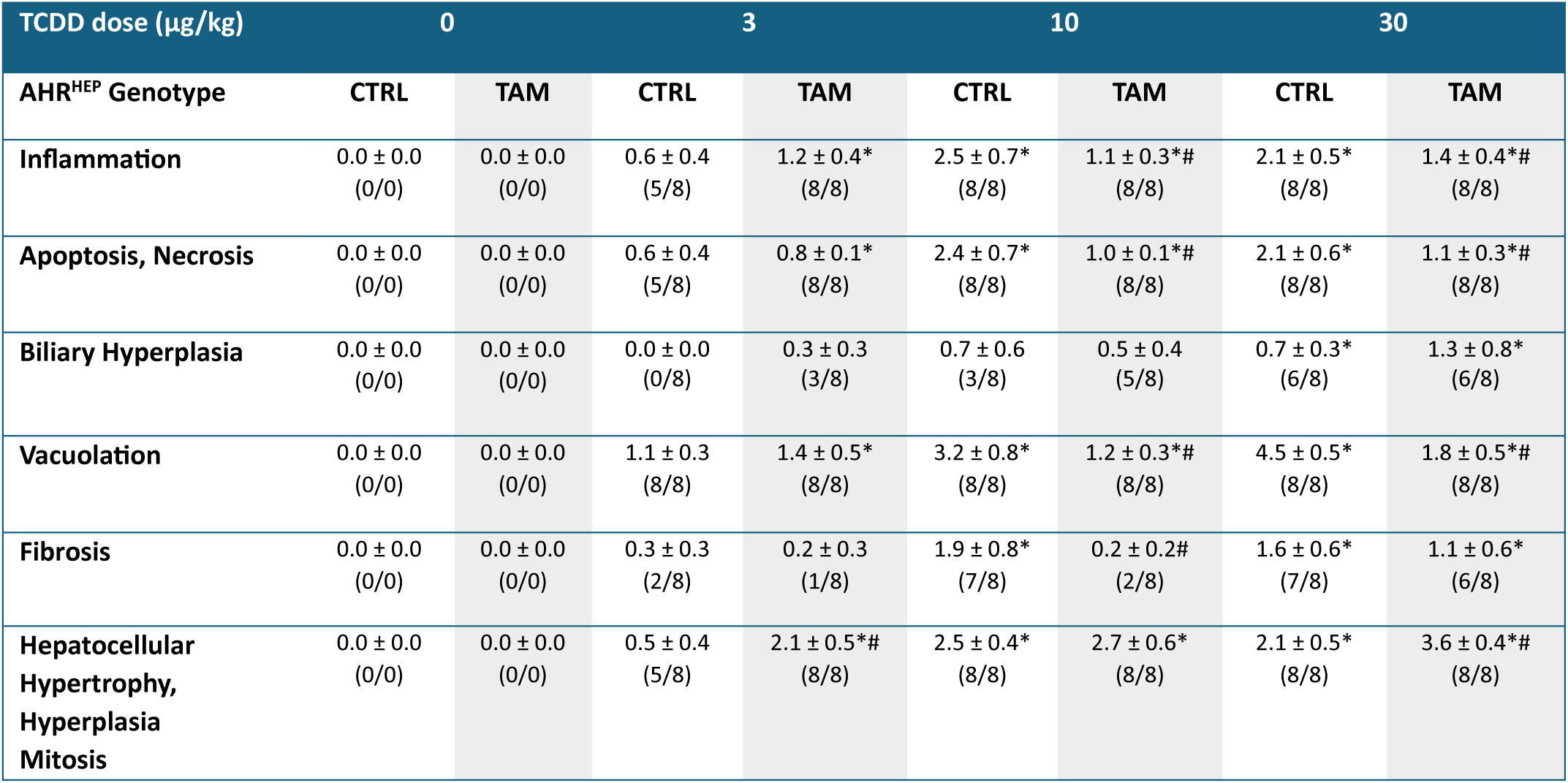
Histopathological assessment of H&E-stained liver sections (n=8).

| **TCDD dose (µg/kg)** | **0** |  | **3** |  | **10** |  | **30** |  |
|---|---|---|---|---|---|---|---|---|
| **AHR<sup>HEP</sup> Genotype** | **CTRL** | **TAM** | **CTRL** | **TAM** | **CTRL** | **TAM** | **CTRL** | **TAM** |
| **Inflammation** | 0.0 ± 0.0<br>(0/0) | 0.0 ± 0.0<br>(0/0) | 0.6 ± 0.4<br>(5/8) | 1.2 ± 0.4*<br>(8/8) | 2.5 ± 0.7*<br>(8/8) | 1.1 ± 0.3*#<br>(8/8) | 2.1 ± 0.5*<br>(8/8) | 1.4 ± 0.4*#<br>(8/8) |
| **Apoptosis, Necrosis** | 0.0 ± 0.0<br>(0/0) | 0.0 ± 0.0<br>(0/0) | 0.6 ± 0.4<br>(5/8) | 0.8 ± 0.1*<br>(8/8) | 2.4 ± 0.7*<br>(8/8) | 1.0 ± 0.1*#<br>(8/8) | 2.1 ± 0.6*<br>(8/8) | 1.1 ± 0.3*#<br>(8/8) |
| **Biliary Hyperplasia** | 0.0 ± 0.0<br>(0/0) | 0.0 ± 0.0<br>(0/0) | 0.0 ± 0.0<br>(0/8) | 0.3 ± 0.3<br>(3/8) | 0.7 ± 0.6<br>(3/8) | 0.5 ± 0.4<br>(5/8) | 0.7 ± 0.3*<br>(6/8) | 1.3 ± 0.8*<br>(6/8) |
| **Vacuolation** | 0.0 ± 0.0<br>(0/0) | 0.0 ± 0.0<br>(0/0) | 1.1 ± 0.3<br>(8/8) | 1.4 ± 0.5*<br>(8/8) | 3.2 ± 0.8*<br>(8/8) | 1.2 ± 0.3*#<br>(8/8) | 4.5 ± 0.5*<br>(8/8) | 1.8 ± 0.5*#<br>(8/8) |
| **Fibrosis** | 0.0 ± 0.0<br>(0/0) | 0.0 ± 0.0<br>(0/0) | 0.3 ± 0.3<br>(2/8) | 0.2 ± 0.3<br>(1/8) | 1.9 ± 0.8*<br>(7/8) | 0.2 ± 0.2#<br>(2/8) | 1.6 ± 0.6*<br>(7/8) | 1.1 ± 0.6*<br>(6/8) |
| **Hepatocellular Hypertrophy, Hyperplasia Mitosis** | 0.0 ± 0.0<br>(0/0) | 0.0 ± 0.0<br>(0/0) | 0.5 ± 0.4<br>(5/8) | 2.1 ± 0.5*#<br>(8/8) | 2.5 ± 0.4*<br>(8/8) | 2.7 ± 0.6*<br>(8/8) | 2.1 ± 0.5*<br>(8/8) | 3.6 ± 0.4*#<br>(8/8) |

### Effects on Steatosis

Oil Red O staining confirmed that the vacuolization in AHR^HEP^*-*CTRL mice induced by TCDD was due to hepatic fat accumulation, while steatosis was essentially absent in TCDD-treated AHR^HEP^*-*TAM mice (**Figure 5A and 5B**). In fact, TCDD decreased hepatic PAS staining, representing glycogen levels, in AHR^HEP^*-*CTRL mice, as previously reported in wild-type C56BL/6Ncrl male mice (*33*) while TCDD had minimal effects on hepatic glycogen levels in AHR^HEP^*-*TAM mice (**Figure 5A and 5B**). Accordingly, *Gys2* and *Gck*, both regulated steps in glycogen biosynthesis, exhibited greater repression in AHR^HEP^*-*CTRL compared to AHR^HEP^*-*TAM mice. There was also AHR genomic enrichment within *Gys2* and *Gck* loci 2 hrs after treatment with 30 µg/kg TCDD in wild-type C57BL/6Ncrl male and female mice in the presence of pDREs (**Supp Fig. S3A**). Collectively, this suggests that the decrease in hepatic glycogen levels was mediated by AHR expression in hepatocytes.

TCDD induced steatosis has been attributed to increased hepatic uptake of dietary and mobilized peripheral fats, the inhibition of lipoprotein assembly and export, and the repression of β-oxidation (*34–36*). Although genes associated with lipid metabolism exhibited comparable differential expression in both models, several were more highly induced in AHR^HEP^*-*CTRL mice (**Supp. Fig. S3**). That included hydrolysis genes, *Lpl* and *Mgll*, as well as the lipid transporter, *Cd36*. In contrast, *Ces3a* which hydrolyzes triglycerides and cholesterol esters, was repressed. *Acot 1*, *2*, *3* and *4*, and *Them7*, which hydrolyze acyl-CoAs to deactivate fatty acids destined for β-oxidation, were also induced in AHR^HEP^*-*CTRL but were largely unaffected by TCDD in AHR^HEP^*-*TAM mice. The higher induction of *Ucp2* in the liver of AHR^HEP^*-*CTRL mice is an adaptive response to excessive fat accumulation that uncouples mitochondrial activity and compromises response to acute energy demands. *Mgll*, *Acot1*, *3* and *4* and *Them7* exhibited negligible induction in AHR^HEP^*-*TAM mice but strong induction in AHR^HEP^*-*CTRL mice with AHR genomic enrichment in the presence of a pDRE suggesting direct AHR regulation. Collectively, these differences are consistent with the development and progression of steatosis to steatohepatitis in AHR^HEP^*-*CTRL and not AHR^HEP^*-*TAM mice.

### Inflammation

F4/80 staining showed increased immune cell infiltration in both models following TCDD treatment (**Table 1**, **Figure 5A**). Although the increase was more pronounced in AHR^HEP^*-*TAM mice, it exhibited a different infiltration pattern compared to AHR^HEP^*-*CTRL mice. Infiltrating immune cells in AHR^HEP^*-*CTRL mice formed more foci while the distribution was spread more uniform throughout the liver section in AHR^HEP^*-*TAM mice which may explain the discrepancy between **Table 1** and **Figure 5B**. RNA-seq also identified increased expression of several inflammation-associated genes in response to TCDD (**Supp. Fig. S4**). Notably, genes associated with macrophage recruitment and injury-associated programs, including *Tlr11*, *Itgam*, *Lyz2*, *Trem2*, *Lgals3*, *Gpnmb* were induced more in AHR^HEP^*-*CTRL mice compared to AHR^HEP^*-*TAM mice. This aligns with the induction of steatosis in AHR^HEP^*-*CTRL mice and suggests that hepatocyte AHR-dependent lipid accumulation may affect macrophage recruitment, activation and programming. Paradoxically, several canonical inflammatory mediators and chemokines, including *Ccl2*, *Ccl3*, *Cxcl1, Cxcl2*, *Tnf*, *Il1b*, and *Lcn2*, were induced in both models, with higher fold changes in AHR^HEP^*-*TAM mice. Only *Tlr11*, *Gpnmb* and *Cxcl1* exhibited AHR genomic recruitment 2 hrs after treatment with 30 µg/kg TCDD in male and female wild-type C57BL/6Ncrl mice suggesting that AHR loss in hepatocytes does not prevent TCDD-induced macrophage recruitment and inflammatory mediator responses. Nevertheless, the histopathology assessment and RNA-seq data indicate that TCDD induced immune cell infiltration in both AHR^HEP^*-*CTRL and AHR^HEP^*-*TAM mice, independent of the severity of steatosis.

### Effects on Fibrosis

Semi-quantitative morphometry found that the incidence and severity of fibrotic changes were also more apparent in AHR^HEP^*-*CTRL mice (**Table 1**). In addition, PSR staining and fibrotic gene expression were higher in AHR^HEP^*-*CTRL mice compared to AHR^HEP^*-*TAM mice (**Figure 5A and 5B, Supp. Fig. S5**). These included genes associated with fibroblast activation (*Vim*, *Acta2*), collagen deposition (*Col1a1, 2; Col3a1; Col4a1, 2, 3, 4, 5, 6*), extracellular matrix remodeling (*Eln*; *Lox*; *Loxl1, 2, 3*) and the development of fibrosis (*Timp1*, *2*; *Serpine1*; *Mmp2, 9, 13*).

### Hepatocellular Hypertrophy and Hyperplasia

Unexpectedly, there was a dramatic increase in hepatocellular hypertrophy and hyperplasia in AHR^HEP^*-*TAM mice treated with TCDD (**Table 1**, **Figure 6 C and D**). Hepatocyte hypertrophy is typically associated with the induction of phase 1 and 2 xenobiotic metabolism. However, TCDD-treated AHR^HEP^*-*TAM mice did not induce phase 1 or 2 genes (**Figure 2D**) yet exhibited marked hepatocyte hypertrophy compared to AHR^HEP^*-*CTRL mice (**Table 1**, **Figure 6C**). The hypertrophic response in AHR^HEP^*-*TAM mice was accompanied by strong evidence of hepatocyte hyperplasia. H&E staining showed hepatocytes from TCDD treated AHR^HEP^*-*TAM mice in various stages of mitosis that occurred randomly across the section suggesting cell division occurred throughout the liver with the typical lobule organization (**Figure 6D**). Ki-67 immunostaining provided additional evidence that hepatocytes from TCDD treated AHR^HEP^*-*TAM mice were experiencing increased cell cycle activity. Increased phospho-histone H3 (P-HH3) staining confirmed the presence of hepatocytes in late G2/M in TCDD treated AHR^HEP^*-*TAM, further demonstrating increased mitotic activity (**Figure 6A and 6B**). As with inflammation and fibrosis, gene expression associated with proliferation was induced in AHR^HEP^*-*CTRL and AHR^HEP^*-*TAM mice (**Supp. Fig. S6**). TCDD elicited comparable gene expression effects on *Hgf*, *Egf* and *Tgfa* while *Hbegf* and *Areg* exhibited higher induction in AHR^HEP^*-*CTRL mice. However, several genes that encoded for immediate early response factors such as *Fos* and *Jun* were more highly induced in AHR^HEP^*-*TAM mice liver samples. Similarly, the pro-proliferative transcription factors (*Foxm1* and *Klf5*) as well as genes that support cell cycle progression, DNA synthesis, chromosome segregation, mitotic regulation, and cellular proliferation (*Cdkn3*, *Ccnb1*, *Ccnb2*, *Cdc20*, *Aurkb*, *Bub1*, *Tk1*) were also more highly induced in AHR^HEP^*-*TAM compared to AHR^HEP^*-*CTRL mice. Several of these including *Jun*, *Foxm1*, *Ccnb1, Aurkb* and *Tk1* exhibited AHR genomic enrichment in wild-type C57BL/6Ncrl mice 2 hrs after treatment with 30 µg/kg TCDD. Together, these results suggest that TCDD increased cell cycle activity in hepatocytes that do not express AHR.

### Hepatocyte AHR ablation enhances TCDD-induced IRE1α/MAPK/AP-1 signaling

We next investigated potential signaling pathways that may contribute to the increased hepatocyte mitotic activity in TCDD-treated AHR^HEP^*-*TAM mice. Functional analysis of RNA-seq data suggested enrichment of differential regulated genes associated with endoplasmic reticulum (ER) stress and mitogen-activated protein kinase (MAPK) signaling (**Figure 7A**). *Ern1*, which encodes for the ER stress sensor, IRE1α, as well as *Map3k5*, were both downregulated by TCDD in AHR^HEP^*-*CTRL mice, whereas there was no repression in AHR^HEP^*-*TAM mice. Both *Map3k5* and *Ern1* exhibited AHR genomic enrichment in the presence of pDREs (**Figure 7B**), suggesting AHR regulation. In contrast, the AP-1 transcription factor components, *Jun* and *Fos,* were induced ∼3-fold but only in TCDD-treated AHR^HEP^*-*TAM mice (**Figure 7B**) while Fos was repressed by TCDD ∼2-fold in AHR^HEP^*-*CTRL mice and exhibited AHR genomic enrichment in the presence of pDREs. In agreement with these transcriptional changes, protein analyses identified differential activation of ER stress and MAPK signaling (**Figure 7C)**. Accordingly, TCDD decreased IRE1α protein levels in AHR^HEP^*-*CTRL mice, whereas TCDD increased phosphorylated IRE1α in AHR^HEP^*-*TAM mice, indicating activation of IRE1α signaling in the absence of hepatocyte AHR following treatment with TCDD. JNK protein levels were also decreased in TCDD-treated AHR^HEP^*-*CTRL mice, while ERK1/2 levels were increased in AHR^HEP^*-*TAM mice. Notably, both total JUN and phosphorylated JUN were increased by TCDD in AHR^HEP^*-*TAM, consistent with enhanced AP-1 signaling. Together, these results strongly suggest that the increase in hepatocyte mitotic activity in AHR^HEP^*-*TAM mice treated with TCDD is associated with the activation of IRE1α/MAPK signaling and increased JUN/AP-1 activity.

**Figure 7.**
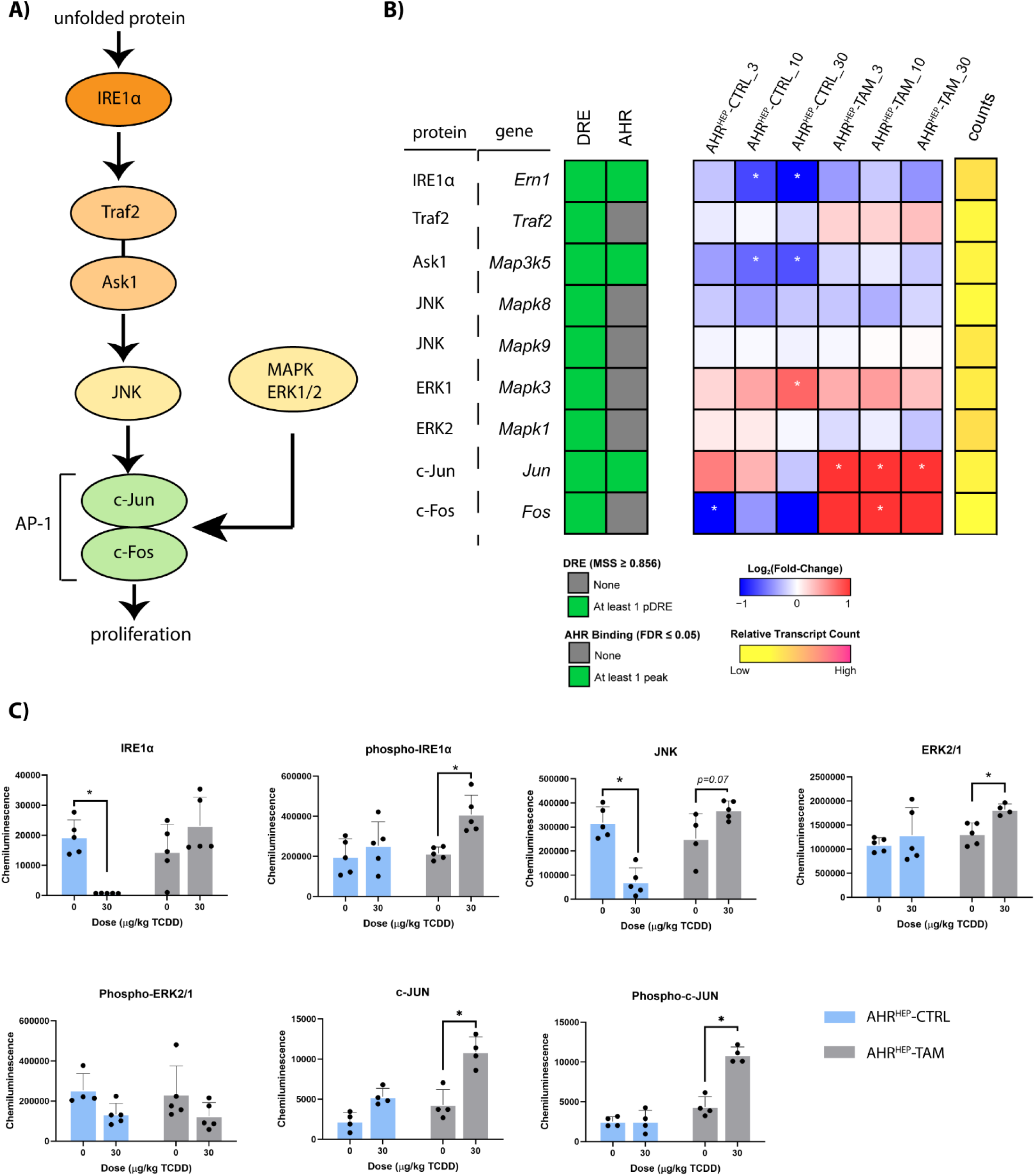
TCDD-Induced IRE1α/MAPK/AP-1 signaling in AHR^HEP^*-*TAM mice. **A)** Proposed mechanism of endoplasmic reticulum (ER) stress induced AP-1-mediated hepatocyte proliferation and/or polyploidization. **B)** Differential expression of genes within the ER stress–AP-1 axis assessed by RNA-Seq. The presence of putative dioxin response elements (pDRE) within the gene loci and *in vivo* hepatic AHR genomic enrichment at 2 hours following treatment with 30 μg/kg TCDD (green). Dose-dependent gene expression was assessed in AHR^HEP^*-*CTRL and AHR^HEP^*-*TAM mice following oral gavage every 4 days for 28 days with vehicle (sesame oil), 3, 10 or 30 µg/kg TCDD. Color scale represents the log2(fold change) in gene expression (n=5). Counts represent the maximum number of raw reads for any treatment group. Low transcript counts (< 100 reads) are denoted in yellow with high transcript counts (> 10,000) denoted in pink. The asterisk (*) indicates differential expression with a Benjamini Hochberg adjusted p-value of 0.05 or less. **C)** Levels of ER stress–AP-1 pathway proteins assessed by capillary electrophoresis using the Wes system (ProteinSimple, San Jose, CA) shown as the mean ± SD (n=4-5). Asterisk (*) indicates p < 0.05 as determined by two-way ANOVA and Tukey’s post-hoc test.

## DISCUSSION

A novel tamoxifen-inducible hepatocyte-specific AHR knockout mouse model that expressed a CRISPR modified *Ahr^dfl/fl^* allele (*Ahr^dV375Afl/fl^*) with high binding affinity for TCDD was used to investigate hepatocyte AHR signaling in response to TCDD, the prototypical and most toxic AHR ligand. AHR hepatocyte ablation was confirmed by loss of CYP1A1 induction in isolated AHR^HEP^*-*TAM primary hepatocytes and liver tissue, while AHR activity was retained in non-parenchymal cells. TCDD induced cytochrome p450 expression, increased ROS levels and oxidative stress as well as markers of liver injury in AHR^HEP^*-*CTRL mice. This was accompanied by hepatic lipid accumulation and increased levels of serum bile acids, hepatic lipid peroxidation and collagen deposition in the liver. These responses were markedly attenuated in TCDD treated AHR^HEP^*-*TAM mice as reflected in the lower induction of fatty acid uptake, lipid metabolism, extracellular matrix remodeling, and fibrogenesis genes. Although hepatocyte AHR deletion did not reduce TCDD-induced immune cell infiltration, the distribution pattern differed between AHR^HEP^*-*CTRL and AHR^HEP^*-*TAM mice with a shift from injury-associated macrophage activation toward enhanced cytokine and chemokine expression. TCDD also induced hepatocyte hypertrophy and hyperplasia in AHR^HEP^*-*TAM mice, as indicated by increased Ki-67 and phospho-histone H3 staining with evidence of hepatocytes undergoing mitosis.

Cytoplasmic vacuolization in hepatocytes due to lipid accumulation is the first histologically identified response to TCDD in wild-type C57/BL/6Ncrl male mice (*37*). TCDD treatment of AHR^HEP^*-*TAM mice clearly shows that AHR expression in hepatocytes is essential for hepatic lipid accumulation. TCDD induces hepatic steatosis by increasing the uptake of mobilized peripheral and dietary fatty acids, repressing β-oxidation, and inhibiting the hepatic export of triglycerides resulting in widespread metabolic reprogramming (*7, 34, 35, 38–41*). Genes involved in fatty acid uptake, intracellular fatty acid transport, lipid hydrolysis, fatty acid activation, acyl-CoA metabolism and mitochondrial metabolism also exhibited negligible differential expression in AHR^HEP^*-*TAM compared to AHR^HEP^*-*CTRL mice following TCDD treatment. In many cases, this could be attributed to the presence of pDREs and AHR genome enrichment suggesting the effects were mediated by the AHR. Overall, these results corroborated with previous conclusions that TCDD promotes steatosis by increasing fatty acid uptake and disrupting hepatic lipid metabolism as opposed to *de novo* lipogenesis (*34, 35, 39*). Furthermore, single nuclei RNA-seq (snRNA-seq) studies show that TCDD disrupted lipid homeostasis not only in hepatocytes but also in non-parenchymal cells. This included the induction of genes involved in lipid transport and steroid biosynthesis, suppression of β-oxidation, and altered linoleic acid metabolism (*42*). However, despite the preservation of AHR signaling in non-parenchymal cells, there was no evidence of TCDD-induced steatosis in AHR^HEP^*-*TAM mice. Therefore, AHR expression in non-parenchymal cells was not sufficient for TCDD to induce hepatic lipid accumulation.

Collagen deposition and the expression of profibrotic genes were also attenuated in AHR^HEP^*-*TAM mice following TCDD treatment. PSR staining similarly revealed less fibrosis in TCDD-treated AHR^HEP^*-*TAM mice. Activation of hepatic stellate cells and extracellular matrix remodeling are downstream consequences of chronic hepatocyte injury, oxidative stress, and inflammation. Accordingly, genes associated with fibroblast activation, extracellular matrix deposition, collagen cross-linking, and matrix remodeling exhibited less induction in AHR^HEP^*-*TAM mice. Our understanding of the role of AHR signaling in liver fibrosis remains incomplete, with its effects depending on the cell type and ligand. AHR activation by TCDD promoted liver fibrosis in mice (*43–45*), whereas whole body AHR null mice developed spontaneous periportal liver fibrosis (*10*). Cell-specific knockout studies suggest this is associated with loss of AHR in HSCs, as fibrosis was observed in HSC-specific AHR knockout mice but not in hepatocyte- or Kupffer cell-specific knockout mice (*46*). Likewise, untreated AHR^HEP^*-*TAM mice did not develop fibrosis within our 28-day study. AHR effects on HSC activation may also be ligand dependent. TCDD-induced fibrosis was also attenuated in AHR^HEP^*-*TAM mice, despite the presence of functional AHR in HSCs. This suggests HSC AHR activation alone is insufficient to drive TCDD induced fibrosis. Alternatively, TCDD may promote fibrosis indirectly due to hepatocyte injury, which then activates HSCs and extracellular matrix remodeling.

Although hepatocyte AHR deletion attenuated steatosis and fibrosis, immune cell infiltration was evident in both models. However, the different transcriptional profiles suggest that the nature of the immune response differed between AHR^HEP^*-*CTRL and AHR^HEP^*-*TAM mice. In AHR^HEP^*-*CTRL mice, TCDD elicited greater induction of macrophage-associated genes consistent with inflammation, clearance of damaged cells, tissue repair and resolution. Specifically, increased expression of *Trem2, Gpnmb, Lgals3*, and *Lyz2* resembles the Lipid-Associated Macrophages (LAMs) or Scar-Associated Macrophages (SAMs) signature, which participate in lipid handling, phagocytosis, and tissue remodeling during liver injury and fibrosis (*47, 48*). These findings are consistent with the steatosis, hepatocellular injury, and fibrosis in TCDD treated AHR^HEP^*-*CTRL mice and suggest recruitment and/or activation of distinct macrophage populations. In contrast, TCDD elicited greater induction of *Tnf, Il1b, Ccl2, Cxcl1,* and *Cxcl2* in AHR^HEP^*-*TAM mice suggesting a proinflammatory signaling response associated with hepatocyte proliferation in the absence of overt liver injury (*49*). Increased *Ccl2* expression is consistent with monocyte recruitment, whereas *Cxcl1* and *Cxcl2* encode chemokines that promote neutrophil recruitment (*50, 51*). TCDD may also directly induce inflammatory cytokines and chemokines through AHR-dependent signaling in nonparenchymal cells, potentially amplifying these responses. Collectively, these findings suggest that hepatocyte AHR signaling influences the nature of the hepatic immune response, with distinct inflammatory and repair-associated programs emerging depending on the existing hepatic environment. Future single-nucleus RNA sequencing studies may help resolve the immune cell subpopulations and transcriptional programs associated with these divergent responses.

The effects of TCDD and AHR signaling on liver growth and hepatocyte proliferation are highly context dependent, with both inhibitory and stimulatory effects reported. AHR deficiency has been associated with increased hepatic proliferation and expression of proliferative markers, as well as enhanced diethylnitrosamine (DEN)-induced liver tumorigenesis (*52*). Conversely, TCDD-activated AHR can exert antiproliferative effects. In mouse Hepa-1 cells, activated AHR interacts with retinoblastoma protein (RB) to repress genes required for S-phase progression (*53–55*). TCDD also suppressed hepatocyte proliferation during liver regeneration by reducing cyclin-dependent kinase 2 (CDK2) activity and inducing the cell-cycle inhibitor p21, thereby impairing cell-cycle progression (*56, 57*). In contrast, liver enlargement induced by the non-toxic AHR ligand, YH439, involved both hepatocyte hypertrophy and proliferation, required AHR expression and was linked to YAP–TEAD signaling (*58*). In our study, TCDD-treated AHR^HEP^*-*TAM mice exhibited hepatocyte enlargement accompanied by increased Ki-67 and phospho-histone H3 (P-HH3) staining. The presence of mitotic figures only in AHR^HEP^*-*TAM mice provided additional evidence of cell cycle activation. Hepatocyte AHR deletion, together with the resulting loss of AHR-dependent CYP450 induction, also affected hepatic sequestration and systemic TCDD distribution, resulting in higher TCDD levels in serum and gWAT. Consequently, hepatic non-parenchymal cells may have experienced different TCDD exposures in AHR^HEP^*-*TAM mice compared to AHR^HEP^*-*CTRL mice. Differences in TCDD tissue distribution, hepatocyte AHR signaling, and liver injury may alter the hepatic microenvironment, including immune-cell recruitment, programming and intercellular signaling (*59–61*). These differences may contribute to the divergent responses observed between the two genotypes, including differences in immune cell responses and the unexpected hepatocyte hypertrophy and hyperplasia in TCDD-treated AHR^HEP^*-*TAM mice.

The liver is organized into repeating functional units, referred to as lobules, whose coordinated organization supports optimal hepatic function. The number of lobules and their growth is tightly regulated to ensure an appropriate liver: body size relationship (*62*). In mice, the adult liver-to-body weight ratio, or liver size set point, is established by ∼PND28, although absolute liver mass continues to increase to ∼PND56. Growth involves both enlargement of existing postnatal lobules and the development of nascent lobules at the periphery, with peripheral vascularization influencing the final adult liver size (*63*). Adult mouse hepatocytes are quiescent with a 200–300-day turnover rate. Nevertheless, they can proliferate rapidly in response to liver injury. Approximately 90% of the mouse hepatocytes are polyploid that can serve as a reservoir for liver regeneration by entering mitosis and replacing damaged cells following exposure to toxic compounds (*64, 65*). Induction of hepatocyte polyploidization has also been attributed to increased liver size with or without cytokinesis depending on the xenobiotic, dose and frequency of exposure (*66*).

Treatment of AHR^HEP^*-*TAM mice with TCDD unexpectedly increased both liver mass and the liver-to-body weight ratio, with evidence of hepatocellular hypertrophy and hyperplasia. Enlarged hepatocytes were observed across all lobular zones with compelling evidence of cell cycle activity. However, in contrast to the enzyme induction commonly associated with other xenobiotic receptor-mediated hepatomegaly (*66, 67*), this occurred in the absence of cytochrome P450 induction. Hepatic lipid accumulation was also absent. The levels of glycogen and bile acids, which are reported to induce hepatocyte proliferation (*68*), were comparable between vehicle and TCDD treated AHR^HEP^*-*TAM mice. Consequently, TCDD induced liver weight in AHR^HEP^*-*TAM mice cannot be attributed to cytochrome P450 induction or the accumulation of hepatic fat or glycogen or induced by bile acids. Liver enlargement only occurred after TCDD exposure indicating that AHR deletion in hepatocytes alone was insufficient to produce this phenotype. This is consistent with previous reports suggesting activation of hepatocyte AHR by TCDD restrains the proliferative response in wild-type models. TCDD treated AHR^HEP^*-*TAM mice exhibited evidence of ER stress and MAPK/AP-1 signaling. Specifically, *Ern1*, encoding the ER stress sensor IRE1α and IRE1α protein levels were repressed by TCDD in AHR^HEP^*-*CTRL mice which would compromise hepatocyte proliferation, while phosphorylated IRE1α was increased in TCDD-treated AHR^HEP^*-*TAM mouse liver extracts. These changes were accompanied by altered MAPK signaling and the induction of the AP-1 transcription factors, *Jun* and *Fos*, together with increased JUN protein levels and its phosphorylation. AP-1 regulates genes involved in cell growth and proliferation in response to MAPK signaling (*69*). This suggests that the loss of AHR signaling in hepatocytes, enabled IRE1α/MAPK/AP-1 activation in response to TCDD that promoted hepatocyte hypertrophy and hyperplasia in the AHR^HEP^*-*TAM mouse liver. TCDD may also promote liver weight increases in AHR^HEP^*-*TAM mice through AHR-dependent signaling still present in non-parenchymal cells that may or may not involve cytokinesis and/or the induction of hepatocyte polyploidization. CellChat analysis of snRNA-seq data showed TCDD increased signaling between hepatocytes and nonparenchymal cells (*70*). Further studies are needed to assess ploidy, identify potential proliferative signals, and establish whether AHR activation in non-parenchymal cells is required to determine whether the increase in liver weight is due to cytokinesis and/or hepatocyte hypertrophy involving polyploidization.

Several limitations should be considered when interpreting results from this study. First is the chimeric nature of the CRISPR-modified AHR. Although this facilitated the use of more relevant TCDD doses, it only included substitution of the valine at position 375 in the *Ahr^d^* allele to an alanine, as found in the high TCDD binding affinity *Ahr^b^* allele. Therefore, in addition to the 3 retained amino acid differences in *Ahr^d^*, the *Ahr*^V375A^ allele also possessed the C-terminal 43 amino acid extension present in the *Ahr^b2^*allele (*18*). The significance of these differences has not been thoroughly explored and may confound comparisons to wild-type C57BL/6Ncrl datasets. The inducible hepatocyte-specific AHR knockout model may also introduce unpredictable interactions between TCDD, CreERT2 protein and TAM that may not be fully considered despite the inclusion of appropriate controls. For example, Alb-CreERT2 can be “leaky”, and may be express in cells other than hepatocytes (*71–73*). In addition, co-treatment of AHR^HEP^ with TAM and TCDD did reduce body weight gain more than TCDD alone, possibility due to residual levels of TAM and its metabolites, N-desmethyl-tamoxifen, 4-hydroxytamoxifen and endoxifen that have been detected in serum up to 7 days post injection (*19*). Cre and TAM-activated CreERT2 can also exert toxicity, although this is highly dependent on the level of expression, targeted tissue, age, TAM dose and level of Cre activity. Indirect evidence suggests that substantial Cre activity is well tolerated by the liver (*74, 75*). In our studies, there were no apparent adverse effects on AHR target gene induction or evidence of histological consequences in the liver due to TAM treatment. Lastly, the RNA-seq studies focused only on protein-coding transcripts and did not consider non-coding RNAs (*76, 77*). It also only represents a snapshot in time after a 28-day study limiting investigation into the mechanisms underlying the dose dependent and temporal progression of steatosis to steatohepatitis with fibrosis. Nevertheless, our *Ahr^V375-fl/fl^;Alb-CreERT2* (AHR^HEP^) mouse model provided temporal and targeted control of *Ahr*^dV375A^ deletion to investigate hepatic AHR biology and the hepatotoxicity of TCDD as intended (*18*).

Overall, our results identified the TCDD-induced liver responses dependent on hepatocyte AHR signaling. AHR expression in hepatocytes was required for the development of steatosis and fibrosis. In contrast, both AHR^HEP^-CTRL and AHR^HEP^-TAM mice exhibited immune cell infiltration, although with distinct spatial distributions, suggesting context-dependent programming. Unexpectedly, the loss of hepatocyte AHR expression shifted the response to TCDD to hepatocellular hypertrophy and hyperplasia, indicating that AHR regulates not only TCDD-induced hepatotoxicity but also hepatocyte hypertrophy and hyperplasia. Consistent with these roles, hepatocellular carcinoma patients with high AHR expression were reported to have significantly poorer survival rates following surgical resection compared to low AHR expression patients (*78*). Collectively, these findings highlight the cell- and context-dependent roles of hepatic AHR signaling and support further investigation of AHR-targeted strategies to enhance liver regeneration following hepatectomy or transplantation.

## MATERIALS and METHODS

### Generation of *Ahr^d^*^V375A*fl/fl*^;*Alb-CreERT2* mice

Mice homozygous for floxed *Ahr^d^*^V375A^ (*Ahr^d^*^V375A*fl/fl*^, provided by Dr. Christopher Bradfield (University of Wisconsin-Madison, Madison, WI; also available from Jackson Laboratory strain designation B6.129(Cg)-*Ahr^em1Bra^*/J, Strain #:035734) were backcrossed onto the C57BL/6J background for six generations (Wilson et al., 2021). To enable hepatocyte-specific deletion, we used mice expressing Cre recombinase fused to a mutant estrogen receptor ligand-binding domain that requires activation by TAM for activity (*Alb-CreERT2*; strain B6.Alb-Cre-ERT2, obtained from Dr. Cornelis Elferink, University of Texas Medical Branch, Galveston, TX) (*79*). Hepatocyte-specific *Ahr^dV375Afl/fl^* knockout mice (*Ahr^d^*^V375A*fl/fl*^*;Alb-CreERT2* and subsequently referred to in this manuscript as AHR^HEP^) were generated by crossing male *Alb-CreERT2^+/-^* mice with *Ahr^d^*^V375A*fl/fl*+/+^ females. To induce hepatocyte-targeted AHR deletion, male and female *Ahr^d^*^V375A*fl/*fl^;Alb*-CreERT2* mice received intraperitoneal injections of either corn oil (AHR^HEP^-CTRL) or 75 mg/kg TAM (AHR^HEP^-TAM) for five consecutive days, followed by 5 recovery days to allow TAM clearance prior to challenging with TCDD.

### Animal housing and treatment

Male and female AHR^HEP^-CTRL and AHR^HEP^-TAM mice were housed in Innovive Innocages (San Diego, CA) containing ALPHA-dri bedding (Shepherd Specialty Papers, Chicago, IL). Mice were maintained under controlled conditions (23 °C, 30–40% humidity, 12-hour light/dark cycle), with free access to Harlan Teklad Rodent Diet 8940 (Madison, WI) and water. On postnatal day (PND) 42-44, mice (n=8 per treatment group) were orally gavaged with sesame oil (vehicle, 100 µl), 3, 10, or 30 μg/kg TCDD (in sesame oil, 100 µl) every four days for 28 days (total of 7 administrations), between zeitgeber time (ZT) 0–3. Mice were euthanized using carbon dioxide followed by cardiac puncture, in accordance with institutional animal care and use approval. The TCDD dose levels considered: (i) the lifelong exposure of humans to AHR ligands, (ii) marked species differences in TCDD elimination rates (1–11 years in humans compared with 8–12 days in mice (*80, 81*)), and (iii) the expectation that individuals are exposed to other environmental AHR agonists. This regimen approached steady state TCDD levels and reliably activated AHR-dependent biological and toxic responses. Although both male and female mice were included in the study design, the main figures focus on male results as they exhibited greater sensitivity compared to female mice. All animal procedures were in accordance with the Michigan State University (MSU) Institutional Animal Care and Use Committee (IACUC; PROTO 202400133) and <u>ARRIVE</u> <u>(Animal Research: Reporting of *In Vivo* Experiments) guidelines</u>.

### Liver histology

Liver sections were fixed in 10% neutral buffered formalin (Sigma-Aldrich, St. Louis, MO), embedded with paraffin, and stained with hematoxylin and eosin (H&E) for general morphometric analysis. For assessment of lipid accumulation previously frozen liver tissues were embedded in optimal cutting temperature (OCT) compound and frozen in 2-methylbutane cooled with liquid nitrogen. Frozen tissues were sectioned at 8 µm, mounted on charged slides, and air-dried for several hours. Sections were fixed in 10% neutral buffered formalin for 5 min, rinsed with running tap water and distilled water for 5 min each, and incubated in 100% propylene glycol for 5 min with agitation. Sections were then stained with working Oil Red O solution at 56°C for 10 min, with agitation after the first 5 min. Slides were rinsed with 80% propylene glycol, washed and counterstain with Gill 2 hematoxylin.

PicroSirius Red (PSR) was performed to verify collagen deposition and F4/80 to determine inflammation based on protocols available online (https://ihcworld.com/2024/01/26/sirius-red-staining-protocol-for-collagen/ and https://www.bio-rad-antibodies.com/indirect-immunostaining-of-paraffin-embedded-tissue-sections.html). F4/80 staining used the following antibodies: Monoclonal Rat anti-Mouse F4/80 Clone: CI:A3-1 – IgG2b (MCA497G) and Goat anti-Rat IgG2b antibody (STAR114P). All processing was performed by the Michigan State University Investigative HistoPathology Laboratory (humanpathology.msu.edu/histology). H&E-stained liver sections were semi-quantitatively assessed by a board-certified veterinary pathologist (JRH), without knowledge of individual animal treatments (*blinded* analysis), using the following criteria: 0, normal (no histopathology); 1, minimal (<10% of liver affected); 2, mild (10-<25% of liver affected): 3, moderate (25-<50% of liver affected); 4, marked (50-<75% of liver affected); 5, severe (75-100% of liver affected). Quantitation of Oil Red O, PAS, F4/80, and PSR staining intensity used QuPath software (0.6.0). The pixel classifier was trained to classify two image features: the staining of interest (lipid accumulation, immune cell infiltration, or fibrosis) and annotation of the entire liver tissue section. The positively stained area was quantified and normalized to the total tissue section area.

### Determination of TCDD concentrations in serum, adipose tissue, and liver

#### Sample extraction and cleanup

Concentrations of 2,3,7,8-tetrachlorodibenzo-*p*-dioxin (TCDD) in serum, gonadal white adipose tissue (gWAT), and liver were determined by Wellington Laboratories Inc. (Guelph, Ontario, Canada). Samples were spiked with ^13^C-labeled 2,3,7,8-TCDD surrogate standard prior to extraction. Approximately 300 mg of liver, 100 mg of adipose tissue, or 60 µL of serum was digested with hydrochloric acid (HCl; 3 mL for liver, 2 mL for adipose tissue, and 1 mL for serum) at room temperature for approximately 60–70 h, 70 h, and 48 h, respectively. Following digestion, TCDD was extracted three times with hexane. The combined extracts were purified by sulfuric acid treatment, multilayer silica chromatography (silica, NaOH/silica, and H₂SO₄/silica), and PX21 carbon chromatography. Adipose tissue extracts underwent an additional lipid freeze-out step before silica column cleanup. Purified extracts were evaporated to dryness and reconstituted in toluene and ^13^C-labelled 1,2,3,4-TCDD injection standard prior to instrumental analysis.

#### GC–MS/MS analysis

The tissue extracts were analyzed using an Agilent 8890 gas chromatograph coupled to a Waters Xevo TQ-XS triple-quadrupole mass spectrometer (HRGC–MS/MS) equipped with an atmospheric-pressure ionization source. The target analytes were chromatographically separated using an HP-5MS UI capillary column (60 m × 0.25 mm × 0.25 µm) and detected using multiple reaction monitoring (MRM), with two transitions monitored per analyte. 2,3,7,8-TCDD was quantified by isotope dilution using multi-point calibration standards containing native and ¹³C-labeled internal standards. Laboratory blanks, matrix blanks, and surrogate recovery measurements were used to assess method performance. Acceptable ^13^C-2,3,7,8-TCDD recoveries ranged from 40% to 120%. TCDD concentrations were reported as pg/g wet weight.

### Immunohistochemistry (IHC)

IHC was performed as previously published with slight modifications (*82*). Formalin-fixed liver sections were de-paraffinized and rehydrated using a regimen of xylene, ethanol, and PBS. After antigen retrieval with IHC buffer (10 mM Tris, 1 mM EDTA, 0.05% Tween-20, pH 9) and heating, slides were cooled and washed with PBS. Endogenous peroxidase was then quenched with 1% H_2_O_2_, and non-target proteins were blocked with the Vectastain ABC Elite kit (PK-6101, Vector Laboratories, Newark, CA). Sections were incubated overnight at 4°C with primary antibodies: CYP1A1 (CRC4, a gift from Dr. Charles D. Rice, Clemson University), Ki67 (12202S; Cell Signaling) or phospho-histone-H3 (9701S; Cell Signaling). Slides were then incubated with secondary antibody (Vector ABC Elite kit) and visualized using ImmPACT NovaRED (SK-4805, Vector Laboratories) with hematoxylin as a counterstain.

### Primary hepatocytes isolation and culture

Primary hepatocytes were isolated from AHR^HEP^-CTRL and AHR^HEP^-TAM mice following perfusion and collagenase digestion (*83*). In brief, livers were perfused with calcium and magnesium-free Hanks’ balanced salt solution (Sigma-Aldrich) supplemented with 0.5 mM EGTA, 5.5 mM glucose, and penicillin-streptomycin (Sigma-Aldrich) followed by perfusion with salt solution supplemented with 1.5 mM calcium chloride, 5.5 mM glucose, penicillin-streptomycin, and 0.02 g of type IV collagenase (Sigma-Aldrich). The liver was removed and gently broken apart with forceps in 10 ml of William’s medium E (WME; Invitrogen, Carlsbad, CA) containing 10% FBS. Clumps and debris were filtered, and the resulting suspension was centrifuged. The pellet was washed with WME and viability was determined by trypan blue exclusion. Hepatocytes with ≥90% viability were plated at 10^6^ cells/well in precoated type I collagen 6-well tissue culture plates containing WME with 5% fetal bovine serum (FBS), 1% penicillin/streptomycin, 1% Glutamax, 0.3 mM ascorbic acid, and 100 nM dexamethasone (*84*). Non-adherent cells were removed after 3 hours, and the WME media (1% ITS Premix Universal Culture Supplement, 1% penicillin/streptomycin, 1% Glutamax, 0.3 mM ascorbic acid, and 100 nM dexamethasone) containing TCDD or DMSO vehicle was added. Cells were then incubated at 37 °C at 5% CO_2_ for 24 hours.

### RNA isolation and qRT-PCR analysis

TRizol (1 mL) was added to each well or to 100 mg of liver tissue. RNA was extracted using an additional 5:1 phenol:chloroform step (Sigma Aldrich) with quantity and purity (260/280 ratio) assessed using a NanoDrop 1000. Conventional PCR was performed using AHR-specific primers (Forward: 5’ CGCAAGCCGGTGCAGAAAAC 3’, Reverse: 5’ ATGGAGGGTGGCTGAAGTGGAGTA 3’) (*79*). Amplicons were separated on a 1.5% agarose gel supplemented with SYBR Safe DNA Gel Stain (Thermo Fisher Scientific, Waltham, MA). Gels were run in 1× TAE buffer at 90–110 V and imaged using a blue-light transilluminator. Total RNA was converted to cDNA with SuperScript II (Invitrogen) using an oligo dT primer. qRT-PCR was performed using iQ SYBR Green Supermix (BioRad, Hercules, CA) on a Bio-Rad CFX Connect Real-Time PCR Detection System. Gene expression relative to vehicle controls was calculated using the 2^−ΔΔCT^ method, where each sample was normalized to the geometric mean of 3 housekeeping genes (*Actb*, *Gapdh*, and *Hprt*). SYBR green Master Mix (Life Technologies, Carlsbad, CA) was used to analyze relative gene expression.

### RNA-Seq analysis

Bulk RNA sequencing was performed on liver samples from AHR^HEP^-CTRL and AHR^HEP^-TAM mice treated with sesame oil vehicle, 3, 10 or 30 µg/kg TCDD (n = 5 biological replicates per treatment group). Total RNA was extracted from frozen liver samples using the TRIzol method with tissue homogenization performed on a BeadBlaster system (Benchmark Scientific, Inc., Sayreville, NJ, USA). RNA quantity and integrity were assessed using an Agilent TapeStation system (Agilent, Santa Clara, CA, USA). RNA sequencing libraries were prepared by Novogene using poly(A) mRNA enrichment and sequenced on an Illumina NovaSeq 6000 to generate 150 bp paired-end reads (Novogene, Beijing, China). Raw sequencing reads were assessed for quality using FastQC v0.12.1. Adapter sequences and low-quality bases were removed using Trimmomatic v0.39. Reads were aligned to the mouse reference genome mm10 using STAR v2.7.1 with default parameters. Transcript counts were generated from aligned reads and used for downstream analyses. Differential expression analysis was conducted in R using DESeq2 v1.34.0. Within each TAM condition, 0 and 75 mg/kg, vehicle-treated samples were compared independently to each TCDD dose group. Genes with an absolute fold-change of 1.5 and a Benjamini Hochberg adjusted p-value of ≥0.05 were considered differentially expressed. Raw and processed RNA sequencing data (GSE354161) are available in the Gene Expression Omnibus. AhR ChIPseq (GSE97634) and computationally identified putative dioxin response elements (pDREs, https://doi.org/10.7910/DVN/JASCVZ) data were previously published (*85*). pDRE data was integrated with *in vivo* hepatic AHR genomic enrichment assessed at 2 hours following oral gavage of male C57BL/6NCrl mice with 30 μg/kg TCDD. Significant AhR ChIPseq binding used a false discovery rate (FDR) ≤ 0.05. pDREs were considered functional with a matrix similarity score (MSS) ≥ 0.856.

### Protein Extraction and Quantitation

Liver samples were homogenized in radioimmunoprecipitation assay (RIPA) buffer supplemented with protease inhibitor cocktail (Sigma-Aldrich). After centrifugation, the total protein concentration was quantified by bicinchoninic acid assay (Sigma-Aldrich). Capillary electrophoresis via the Wes system (ProteinSimple, San Jose, CA) was used to detect IRE1α (#3294; 1:50; Cell Signaling, Danvers, MA), Erk1/Erk2 (#4695; 1:50; Cell Signaling), phospho-Erk1/Erk2 (#9101; 1:50; Cell Signaling), JNK (#51151-1-AP; 1:50; Proteintech, Rosemont, IL) and phospho-IRE1α (#NB100-2323; 1:50; Novus Biologicals, Centennial, CO). Chemiluminescence signals were exported from Compass software (ProteinSimple) and used to calculate fold-change.

### Western blot

Protein lysates (20 µg) were separated by sodium dodecyl sulfate-polyacrylamide gel electrophoresis (SDS-PAGE) in 10% gels (Bio-Rad) and transferred to nitrocellulose membranes (GE Healthcare, Chicago, IL) by wet-transfer system (Mini Trans-Blot Cell Unit, BioRad) operated at 100V for 45 min. The membranes were blocked with 5% nonfat milk (in Tris-buffered saline [TBS] + 0.01% Tween) for 1 h and incubated overnight at 4 °C with primary antibodies targeting: c-Jun (#9165; 1:1000; Cell Signaling), phospho-c-Jun (#3270; 1:1000; Cell Signaling), or β-actin (#4970s; 1:1000; Cell Signaling). The next day, membranes were washed with 0.1% TBST at room temperature and probed with horseradish peroxidase (HRP)-linked secondary antibodies (1:3000; 7074S; Cell Signaling). Detection was performed using ECL kit (Millipore Corporation, Billerica, MA) and scanned on a Sapphire Biomolecular Imager (Azure Biosystem, Dublin, CA). Two images were captured for each blot to visualize the chemiluminescence protein recognized by the antibody, and a second to visualize the molecular weight marker. The images were overlayed to estimate the molecular weight of the visible bands.

### Clinical chemistry and colorimetric assays

Serum cholesterol (MAK043; Sigma), triglycerides (Ab65336; Abcam), free fatty acids (ab65341; Abcam), glucose (EIAGLUC; Invitrogen), alanine aminotransferase activity (MAK052; Sigma), lipid peroxidation (ab233471; Abcam) and total bile acids (STA-631; BioLabs, Boston, MA) were quantified using the respective commercial assay kits according to the manufacturers’ instructions.

## Acknowledgments

We thank David Potter of Wellington Laboratories Inc. for assistance with the quantification of TCDD concentrations in mouse tissues.

## Funding

This project was supported by the National Institute of Environmental Health Sciences R01ES029541 and SRP P42ES004911 to TZ. TZ is partially supported by AgBioResearch at Michigan State University. CAB is supported by the National Institute of Environmental Health Sciences R35 ES028377.

## Author contributions

Conceptualization: TZ and RN; Methodology: TZ, RN, RHW, CAB and KO; Funding acquisition: TZ and RN; Investigation: KO, RN, GC, ZS, WS, AW, AV, FU, DV, CR, NS, JR, JS, RHW, CAB, TZ; Software: RN, GC, WS; Supervision: TZ, RN and KO; Visualization: KO and GC; Writing—original draft: KO and TZ; Writing—review &editing: KO, RN, GC, ZS, WS, AW, AV, FU, DV, CR, NS, JR, JS, RHW, CAB, TZ

## Competing interests

The authors declare that they have no competing interests.

## Data availability

All data needed to evaluate and reproduce the conclusions in the paper are present in the paper and/or the Supplementary Materials. RNA-seq data was deposited to GEO under the accession identifier GSE354161.

## Declaration of Generative AI Use

This manuscript was improved for clarity and conciseness with the support of OpenAI’s ChatGPT-4 (OpenAI, San Francisco, CA). The authors carefully reviewed and edited the language model’s output to ensure the accuracy and integrity of the content. The authors assume full responsibility for the final version of the manuscript.

## Supplementary Materials

This manuscript contains supplementary materials: Supplementary Figure S1-S7 and Supplementary Table 1.

## Notes

### Competing Interest Statement

The authors have declared no competing interest.

