## Supplementary_Materials for "Inducible Hepatocyte Aryl Hydrocarbon Receptor Ablation Prevented TCDD-induced Steatosis and Fibrosis but Enhanced Hyperplasia and Hypertrophy"

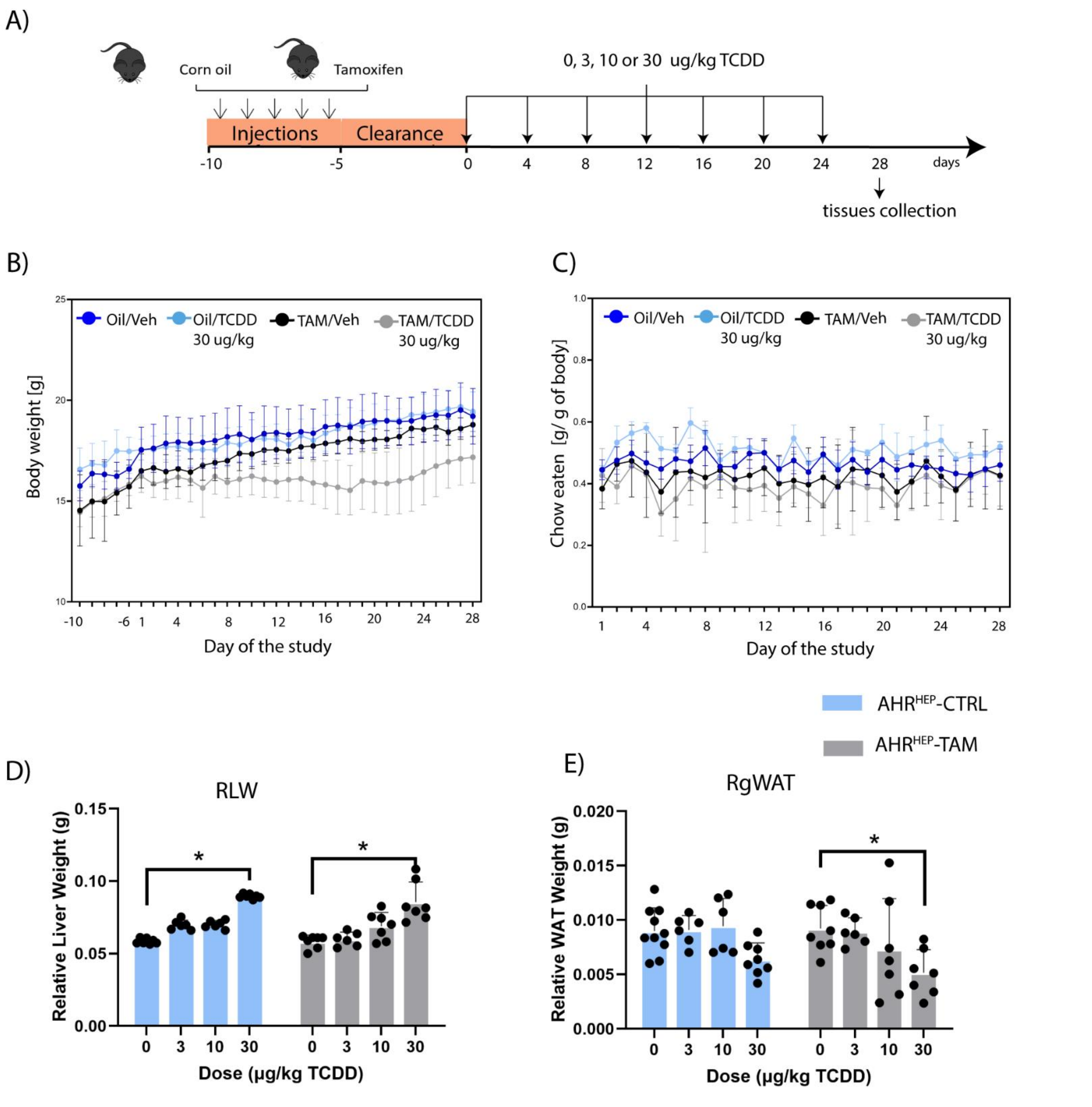


**Supplementary Figure S1.** Experimental design and physiological responses to TCDD in female AHR$^{HEP}$-CTRL and AHR$^{HEP}$-TAM mice. **A)** Schematic representation of experimental design. Hepatocyte-specific AHR ablation was achieved by intraperitoneally injecting AHR$^{HEP}$ mice with 75 mg/kg tamoxifen (TAM) for five consecutive days, followed by 5 recovery days to allow TAM clearance. Male and female mice were then orally gavaged with sesame oil vehicle or TCDD every 4 day for 28 days. **B)** Body weight assessment in female mice. **C)** Average chow consumption per treatment group presented as g of food per g of body weight in female mice. **D)** Relative liver weight (RLW) in female mice and **E)** relative gonadal white adipose tissue (RgWAT) weight in female mice on day 28, 4 days after the last oral gavage of TCDD. Bars represent mean ± SD. Asterisk (*) indicates difference ($p < 0.05$) determined by two-way ANOVA and Tukey's post-hoc test.

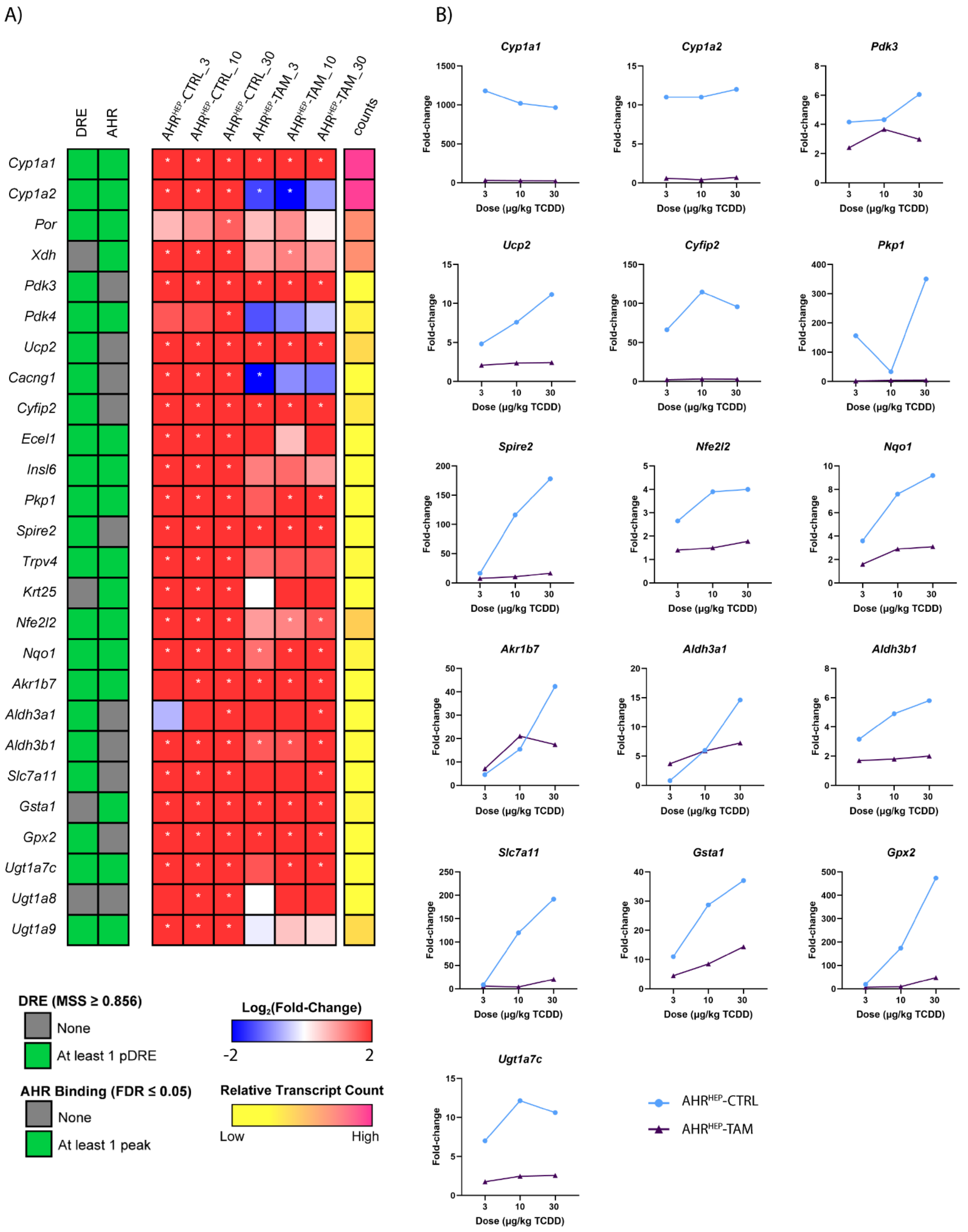
A)
DRE
AHR
AHRHEP-CTRL_3
AHRHEP-CTRL_10
AHRHEP-CTRL_30
AHRHEP-TAM_3
AHRHEP-TAM_10
AHRHEP-TAM_30
counts
Cyp1a1
Cyp1a2
Por
Xdh
Pdk3
Pdk4
Ucp2
Cacng1
Cyfip2
Ecel1
Insl6
Pkp1
Spire2
Trpv4
Krt25
Nfe2l2
Nqo1
Akr1b7
Aldh3a1
Aldh3b1
Slc7a11
Gsta1
Gpx2
Ugt1a7c
Ugt1a8
Ugt1a9
DRE (MSS ≥ 0.856)
None
At least 1 pDRE
AHR Binding (FDR ≤ 0.05)
None
At least 1 peak
Log2(Fold-Change)
-2
2
Relative Transcript Count
Low
High
B)
Cyp1a1
Cyp1a2
Pdk3
Ucp2
Cyfip2
Pkp1
Spire2
Nfe2l2
Nqo1
Akr1b7
Aldh3a1
Aldh3b1
Slc7a11
Gsta1
Gpx2
Ugt1a7c
Fold-change
Dose (μg/kg TCDD)
AHRHEP-CTRL
AHRHEP-TAM

**Supplementary Figure S2. Dose dependent effect of TCDD on gene expression associated with cellular stress and oxidative stress responses in AHR$^{HEP}$-CTRL and AHR$^{HEP}$-TAM male mice.** Mice (n=5) were orally gavaged every 4 days for 28 days with vehicle (sesame oil), 3, 10 or 30 µg/kg TCDD. Hepatic gene expression was assessed by RNA-Seq. **(A)** Heatmap of hepatic cellular stress and oxidative stress genes with AHR regulatory annotation. The presence of putative dioxin response elements (pDRE) within the gene locus and *in vivo* hepatic AHR genomic enrichment at 2 hours in male wild type C57BL/6Ncrl mice following oral gavage with 30 µg/kg TCDD (green). Color scale represents the log2(fold change) for differential gene expression determined by RNA-Seq analysis (n=5). Counts represent the maximum number of raw reads for any treatment group. Low transcript counts (< 100 reads) are denoted in yellow with high transcript counts (> 10,000) denoted in pink. The asterisk (*) indicates differential expression with a Benjamini Hochberg adjusted p-value of ≥0.05. **(B)** Dose response profiles of selected cellular stress and oxidative stress responses-associated genes. Line graphs show the differences in the level of induction for representative genes, highlighting greater level of induction in AHR$^{HEP}$-CTRL mice compared to AHR$^{HEP}$-TAM male mice.

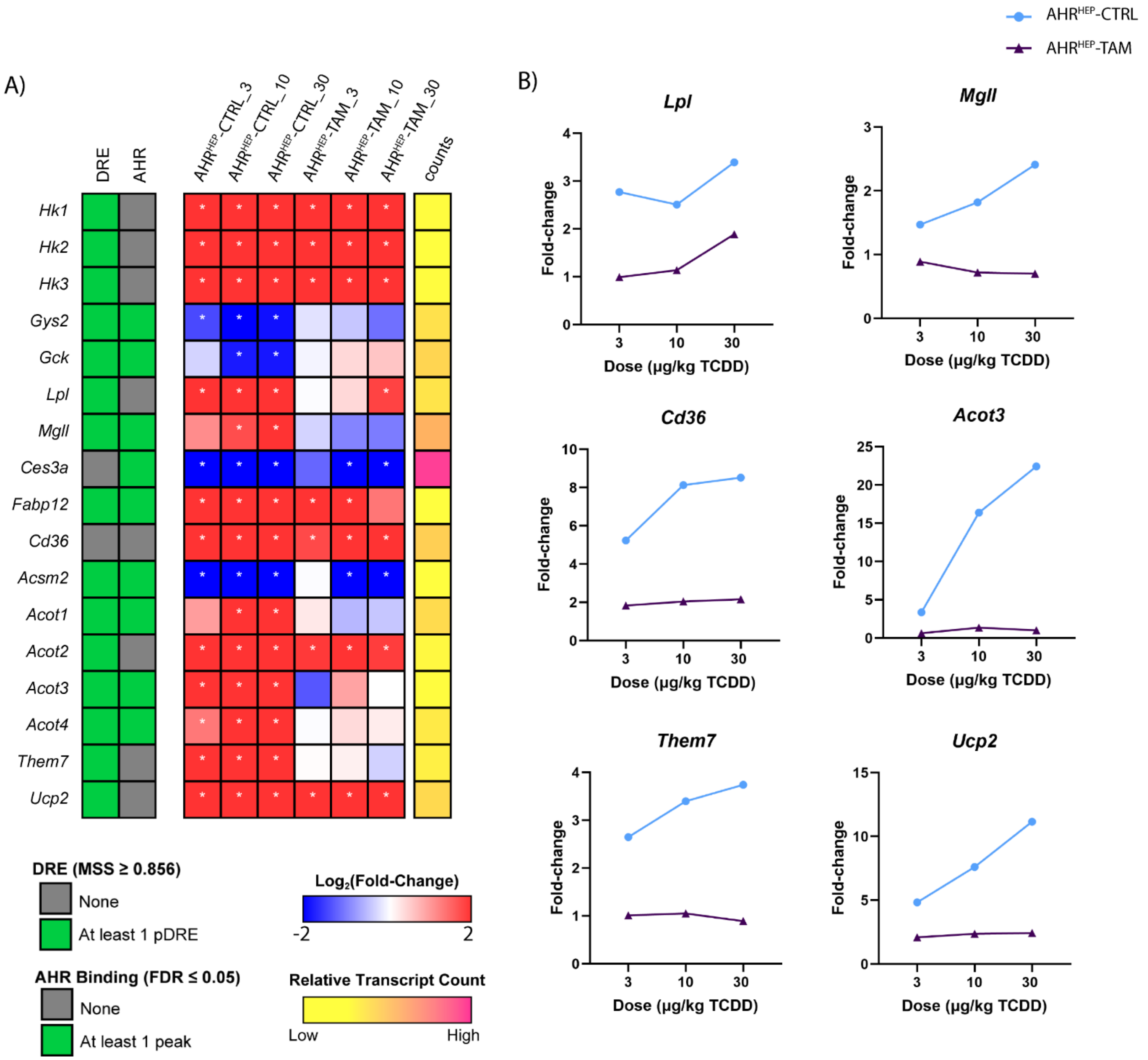


**Supplementary Figure S3. Dose dependent effect of TCDD on the expression of genes associated with glucose metabolism and hepatic steatosis in AHR$^{HEP}$-CTRL and AHR$^{HEP}$-TAM male mice.** Mice (n=5) were orally gavaged every 4 days for 28 days with vehicle (sesame oil), 3, 10 or 30 µg/kg TCDD. Hepatic gene expression was assessed by RNA-Seq. **(A)** Heatmap of differential gene expression associated with steatosis. The presence of putative dioxin response elements (pDRE) within the gene locus and *in vivo* hepatic AHR genomic enrichment at 2 hours in wild type C57BL/6Ncrl male mice following treatment with 30 µg/kg TCDD (green). Color scale represents the log2(fold change) for differential gene expression determined by RNA-Seq analysis (n=5). Counts represent the maximum number of raw reads for any treatment group. Low transcript counts (< 100 reads) are denoted in yellow with high transcript counts (> 10,000) denoted in pink. The asterisk (*) indicates differential expression based on a Benjamini Hochberg adjusted p-value 0.05 or less. **(B)** Dose response expression profiles of selected steatosis-associated genes. Line graphs show the fold change differences for representative genes, highlighting the greater level of induction in AHR$^{HEP}$-CTRL mice compared to AHR$^{HEP}$-TAM male mice.

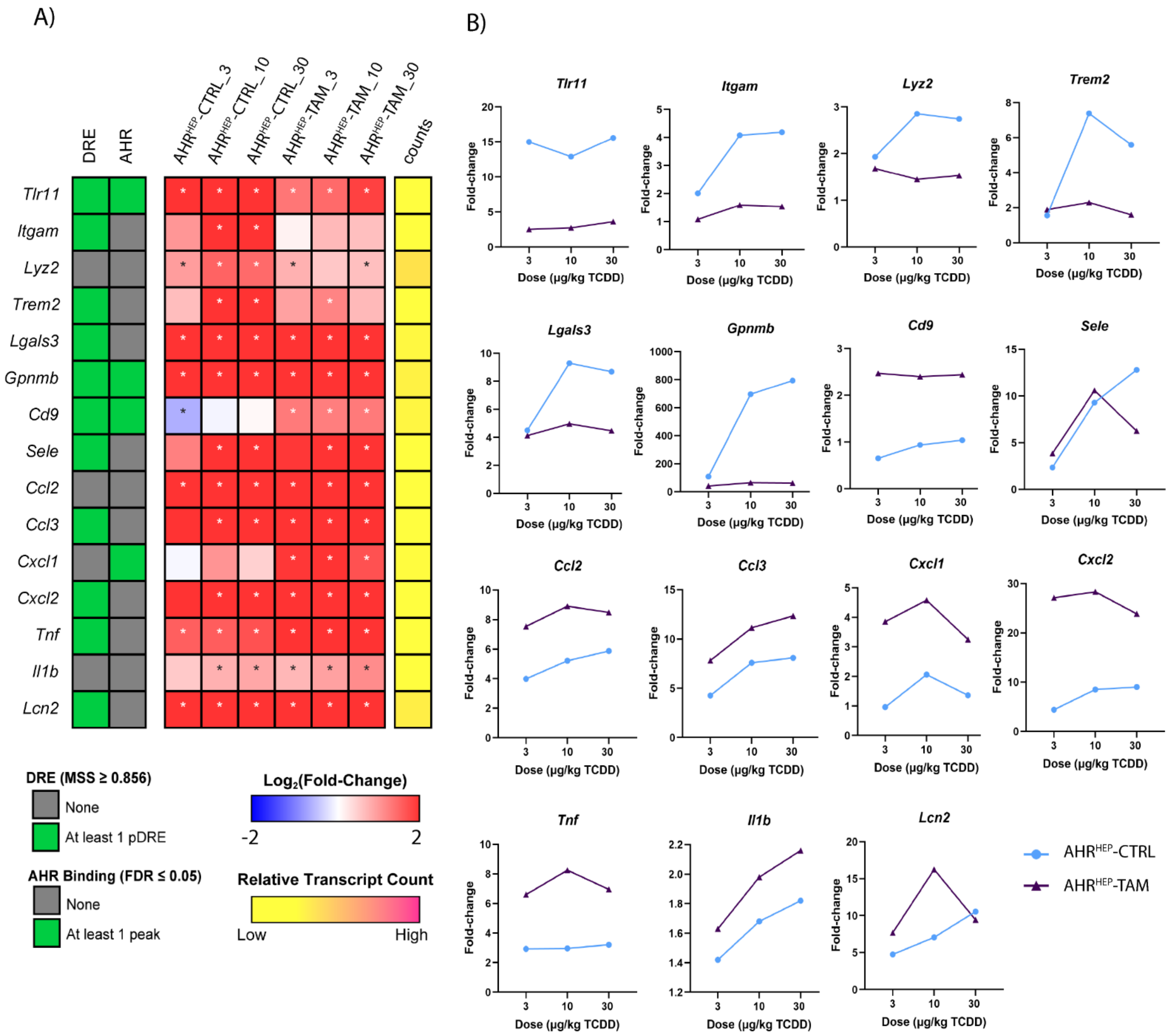


**Supplementary Figure S4. Dose dependent effect of TCDD on the expression of genes associated with hepatic immune cell infiltration in AHR$^{HEP}$-CTRL and AHR$^{HEP}$-TAM male mice.** Mice (n=5) were orally gavaged every 4 days for 28 days with vehicle (sesame oil), 3, 10 or 30 µg/kg TCDD. Hepatic gene expression was assessed by RNA-Seq. **(A)** Heatmap of hepatic inflammation genes with AHR regulatory annotation. The presence of putative dioxin response elements (pDRE) within the gene locus and *in vivo* hepatic AHR genomic enrichment at 2 hours in male wild type C57BL/6Ncrl mice following oral gavage with 30 µg/kg TCDD (green). Color scale represents the log2(fold change) for differential gene expression determined by RNA-Seq analysis (n=5). Counts represent the maximum number of raw reads for any treatment group. Low transcript counts (< 100 reads) are denoted in yellow with high transcript counts (> 10,000) denoted in pink. The asterisk (*) indicates differential expression with a Benjamini Hochberg adjusted p-value of ≥0.05. **(B)** Dose response profiles of selected inflammation-associated genes. Line graphs show the fold-change differences for representative genes associated with hepatic immune cell infiltration in $AHR^{HEP}$-CTRL and $AHR^{HEP}$-TAM male mice.

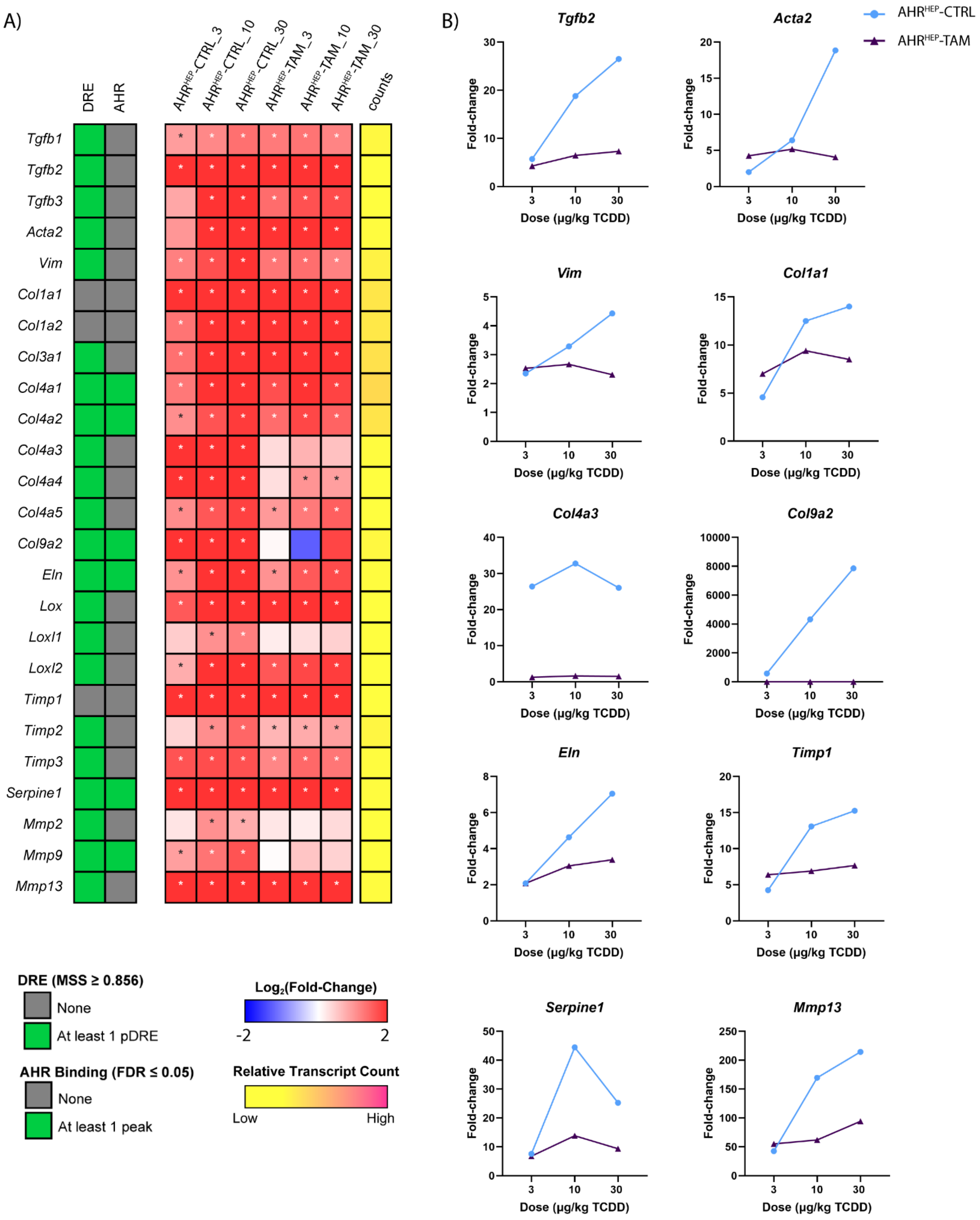

A)
DRE
AHR
AHR^HEP-CTRL_3
AHR^HEP-CTRL_10
AHR^HEP-CTRL_30
AHR^HEP-TAM_3
AHR^HEP-TAM_10
AHR^HEP-TAM_30
counts
Tgfb1
Tgfb2
Tgfb3
Acta2
Vim
Col1a1
Col1a2
Col3a1
Col4a1
Col4a2
Col4a3
Col4a4
Col4a5
Col9a2
Eln
Lox
Loxl1
Loxl2
Timp1
Timp2
Timp3
Serpine1
Mmp2
Mmp9
Mmp13
DRE (MSS ≥ 0.856)
None
At least 1 pDRE
AHR Binding (FDR ≤ 0.05)
None
At least 1 peak
Log2(Fold-Change)
-2
2
Relative Transcript Count
Low
High
B)
AHR^HEP-CTRL
AHR^HEP-TAM
Tgfb2
Acta2
Vim
Col1a1
Col4a3
Col9a2
Eln
Timp1
Serpine1
Mmp13
Fold-change
Dose (µg/kg TCDD)
3
10
30

**Supplementary Figure S5. Dose dependent effect of TCDD on gene expression associated with hepatic fibrosis in AHR$^{HEP}$-CTRL and AHR$^{HEP}$-TAM male mice.** Mice (n=5) were orally gavaged every 4 days for 28 days with vehicle (sesame oil), 3, 10 or 30 µg/kg TCDD. Hepatic gene expression was assessed by RNA-Seq. **(A)** Heatmap of hepatic fibrosis genes with AHR regulatory annotation. The presence of putative dioxin response elements (pDRE) within the gene locus and *in vivo* hepatic AHR genomic enrichment at 2 hours in male wild type C57BL/6Ncrl mice following oral gavage with 30 µg/kg TCDD (green). Color scale represents the log2(fold change) for differential gene expression determined by RNA-Seq analysis (n=5). Counts represent the maximum number of raw reads for any treatment group. Low transcript counts (< 100 reads) are denoted in yellow with high transcript counts (> 10,000) denoted in pink. The asterisk (*) indicates differential expression with a Benjamini Hochberg adjusted p-value of ≥0.05. **(B)** Dose response profiles of selected fibrosis-associated genes. Line graphs show the fold change differences for representative genes, highlighting the greater level of induction of genes associated with fibrosis in AHR$^{HEP}$-CTRL mice compared to AHR$^{HEP}$-TAM male mice.

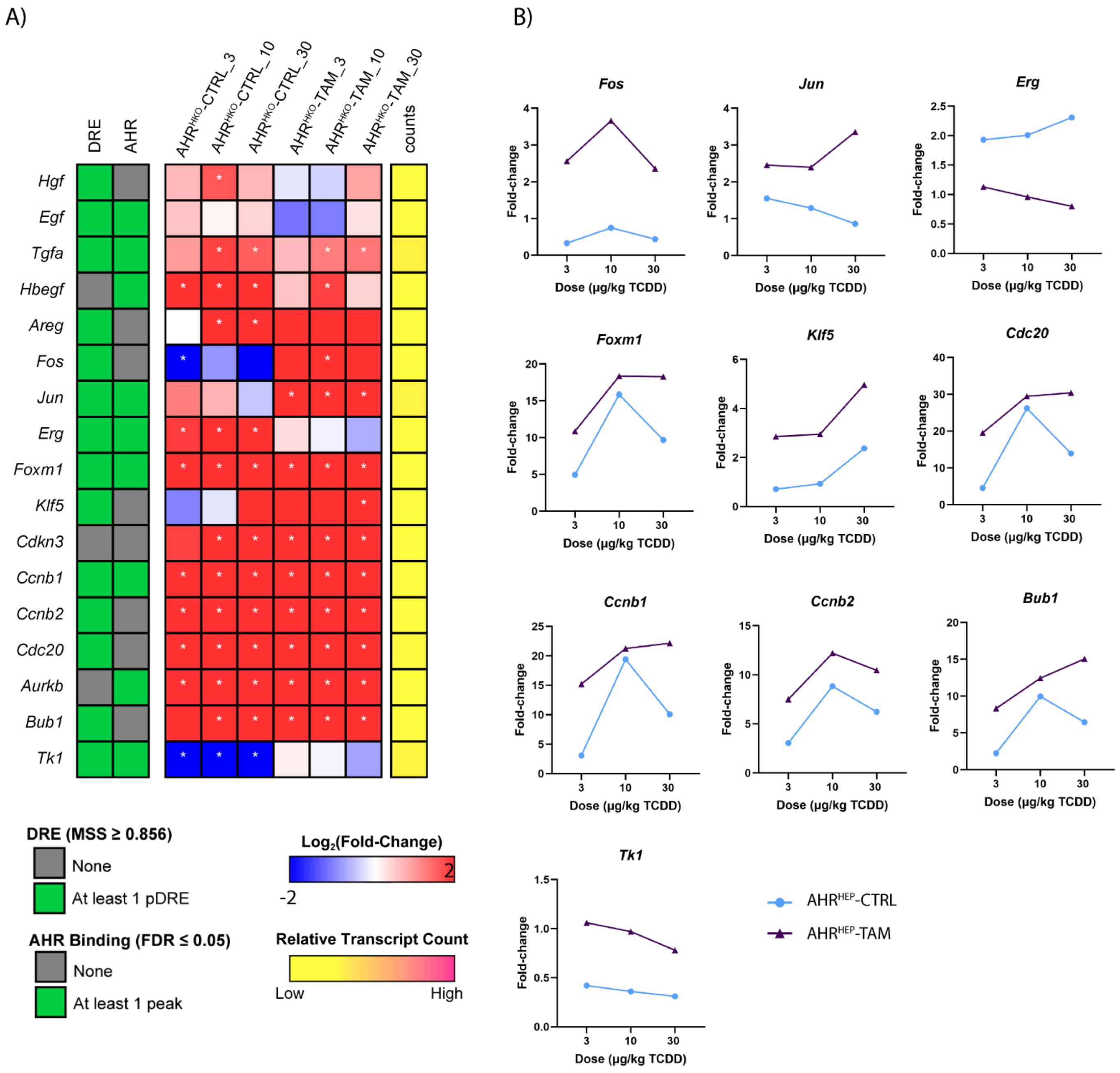


**Supplementary Figure S6. Dose dependent effect of TCDD on the expression of genes associated with cell proliferation in AHR$^{HEP}$-CTRL and AHR$^{HEP}$-TAM male mice.** Mice (n=5) were orally gavaged every 4 days for 28 days with vehicle (sesame oil), 3, 10 or 30 µg/kg TCDD. Hepatic gene expression was assessed by RNA-Seq. **(A)** Heatmap of cell proliferation genes with AHR regulatory annotation. The presence of putative dioxin response elements (pDRE) within the gene locus and *in vivo* hepatic AHR genomic enrichment at 2 hours in male wild type C57BL/6Ncrl mice following oral gavage with 30 µg/kg TCDD (green). Color scale represents the log2(fold change) for differential gene expression determined by RNA-Seq analysis (n=5). Counts represent the maximum number of raw reads for any treatment group. Low transcript counts (< 100 reads) are denoted in yellow with high transcript counts (> 10,000) denoted in pink. The asterisk (*) indicates differential expression with a Benjamini Hochberg adjusted p-value of ≥0.05. **(B)** Dose response profiles of selected proliferation-associated genes. Line graphs show the fold change differences for representative genes, highlighting the greater level of induction of genes associated with proliferation in AHR$^{HEP}$-TAM mice compared to AHR$^{HEP}$-CTRL male mice.

**Supplementary Table S1. Dose-dependent accumulation of TCDD in serum, gonadal white adipose tissue (gWAT), and liver of AHR$^{HEP}$-CTRL and AHR$^{HEP}$-TAM male mice.** TCDD concentrations are expressed in parts per trillion (ppt). Data are presented as mean ± SEM (n = 5 per group).

| TCDD dose (µg/kg) | AHR$^{HEP}$ Genotype | Serum | gWAT | Liver |
|---|---|---|---|---|
| **0** | CTRL | 0.24 ± 0.11 | 89.6 ± 67.7 | 133 ± 60.8 |
| **0** | TAM | 0.51 ± 0.47 | 372 ± 152 | 216 ± 81.3 |
| **3** | CTRL | 46.5 ± 14.1 | 1.23e+04 ± 2.21e+03 | 6.92e+04 ± 1.12e+04 |
| **3** | TAM | 164 ± 86.2 | 5.70e+04 ± 2.73e+04 | 5.76e+03 ± 2.32e+03 |
| **10** | CTRL | 182 ± 38.3 | 5.80e+04 ± 7.93e+03 | 2.35e+05 ± 4.35e+04 |
| **10** | TAM | 1.01e+03 ± 235 | 2.82e+05 ± 6.08e+04 | 2.33e+04 ± 2.6e+03 |
| **30** | CTRL | 458 ± 124 | 1.20e+05 ± 2.06e+04 | 6.07e+05 ± 6.85e+04 |
| **30** | TAM | 1.89e+03 ± 738 | 5.06e+05 ± 1.68e+05 | 7.34e+04 ± 1.65e+04 |
